# Tissue-specific sex difference in mouse eye and brain metabolome under fed and fasted states

**DOI:** 10.1101/2023.01.10.523270

**Authors:** Meghashri Saravanan, Rong Xu, Olivia Roby, Yekai Wang, Siyan Zhu, Amy Lu, Jianhai Du

## Abstract

**Purpose:** Visual physiology and various ocular diseases demonstrate sexual dimorphisms; however, how sex influences metabolism in different eye tissues remains undetermined. This study aims to address common and tissue-specific sex differences in metabolism in the retina, retinal pigment epithelium (RPE), lens and brain under fed and fasted conditions.

**Methods:** After ad libitum fed or deprived of food for 18 hours, mouse eye tissues (retina, RPE/choroid, and lens), brain, and plasma were harvested for targeted metabolomics. The data were analyzed with both Partial least squares-discriminant analysis (PLS-DA) and Volcano Plot analysis.

**Results:** Among 133 metabolites that cover major metabolic pathways, we found 9-45 metabolites that are sex-different in different tissues under the fed state and 6-18 metabolites under the fasted state. Among these sex-different metabolites, 33 were changed in two or more tissues, and 64 were tissue-specific. Pantothenic acid, hypotaurine and 4-hydroxyproline were the top commonly changed metabolites. Lens and retina had the most tissue-specific sex-different metabolites enriched in the metabolism of amino acid, nucleotide, lipids and TCA cycle. Lens and brain had more similar sex-different metabolites than other occular tissues. Female RPE and female brain were more sensitive to fasting with more reduced metabolites in amino acid metabolism, TCA cycle and glycolysis. The plasma had the least sex-different metabolites with very few overlapping changes with tissues.

**Conclusion:** Sex has a strong influence on eye and brain metabolism in tissue-specific and metabolic state-specific manners. Our findings may implicate the sexual dimorphisms in eye physiology and susceptibility to ocular diseases.

## INTRODUCTION

Sexual dimorphisms have been reported in vertebrate eyes, including photoreceptor cell distribution, visual acuity, color perception and disease susceptibility. The males have more relative number of L- and M-cone photoreceptors, thicker macula, stronger response to blue light stimuli, and greater sensitivity for fine detail and rapidly moving stimuli^1-5^. The females have a higher density of lens epithelium and more irons in the retina and retinal pigment epithelium (RPE)^6-8^. Men are known for the high prevalence of color blindness^9^; however, women are more susceptible to age-related macular degeneration (AMD), cataract and glaucoma^10-13^. Although sex differences in eye physiology and pathology are well established, the biochemical basis for these sex differences in different eye tissues remains unknown.

Metabolome is a collection of metabolites such as carbohydrates, amino acids, nucleotides, fatty acids and vitamins in the cell or tissue, serving as substrates, products, cofactors, or ligands for biochemical reactions, nutrient transport and cell signaling^14^. These metabolites are not only the products of metabolic gene and protein expression, but also reflect interactions with the environment such as microbiome, diets and exposure^15-17^. Quantifying metabolites by metabolomics is increasingly important in eye research to identify tissue-specific metabolism in healthy ocular tissues and mechanisms or biomarkers in ocular diseases^18-22^. Notably, the ocular tissues have specialized metabolism, which may underlie various ocular diseases that cause blindness. Like tumors, the neural retina has the Warburg effect or aerobic glycolysis to produce large amounts of lactate from glucose. Many mutations of metabolic genes in glycolysis, tricarboxylic acid (TCA) cycle, and nucleotide metabolism only cause retinal degeneration in humans^23-25^. Retinal pigment epithelium (RPE), a single layer of epithelial cells, resides between the neural retina and choroid circulation. RPE metabolism is critical to the survival of the neural retina. The defects in RPE metabolism are attributed to inherited retinal degeneration and AMD, the leading cause of blindness in the elder population^23, 26, 27^. The lens is a transparent tissue that relies on nutrients, especially glucose, from the aqueous humor through the blood-aqueous barrier^28^. Metabolic disturbance of lens metabolism can cause the loss of transparency or cataract, a common ocular disease in the elderly^29, 30^. Sex differences in glucose, lipid and amino acid metabolism are well-studied in adipose, muscle and liver^31-34^; however, the sex differences in eye metabolism have not been determined and appreciated.

In this study, we used a targeted metabolomics approach to quantify 133 metabolites covering major intermediates in the metabolism of glucose, amino acids, nucleotides, fatty acids and vitamins in the neural retina, RPE and lens from male and female mice under fed or fasted conditions. We also quantified the metabolites from the brain and plasma to identify common and tissue-specific sex differences in metabolism. We have found 97 sex-different metabolites and 64 metabolites show tissue-specific sex differences. Our findings demonstrate strong tissue-specific and sex-specific differences in eye metabolome, and these differences may implicate the sex differences in eye physiology and susceptibility to diseases.

## METHODS

### Animals

12-week-old C57 BL/6J mice of both sexes were purchased from JAX (Stock #:000664). The fed group had ad libitum access to food, but the food was removed for 18 hours after 4 PM in the fasted group. All mouse experiments were performed in accordance with guidelines by the National Institutes of Health and ARVO Statement for the Use of Animals in Ophthalmic and Vision Research, and the protocols were approved by the Institutional Animal Care and Use Committee of West Virginia University.

### Isolation of retina, RPE/Choroid, lens, brain and plasma

All mice were euthanized through quick cervical dislocation. Enucleated eyeballs were cleaned for the lingering fat and muscle tissue and dissected to isolate the retina, RPE/choroid and lens as previously reported^21, 35^. Another technician quickly drew blood from the heart into microtubes with 10 µl of 0.5 mM EDTA and centrifuged at 3000 rpm at 4°C for 15 min to collect the supernatant to fresh microtubes. The whole brain tissue was rapidly dissected and snap-frozen in liquid nitrogen. All the harvested samples were stored at -80°C before use.

### Metabolite extraction and preparation

Metabolites from the retina, RPE/choroid, lens and brain were extracted with 80% cold menthol together with internal stand norvaline (1mM) as described^36, 37^. Plasma metabolites were extracted by mixing 10 µl of plasma with 40 µl of cold methanol with norvaline. The mix was centrifuged and 10 µl of supernatant was used for metabolite analysis. Protein pellets from the extraction were measured for protein concentrations using BCA assay for normalization (ThermoFisher, # 23227)^38^. All the metabolite extracts were dried before targeted metabolomics.

### Targeted metabolomics

Targeted metabolomics was performed as described in detail before with Liquid chromatography-mass spectrometry (LC MS/MS) and gas chromatography-mass spectrometry (GC MS)^22, 37^. A total of 133 metabolites that cover major metabolic pathways were quantified (see detailed pathways and parameters in **Supplemental Table S1**). A Shimadzu LC Nexera X2 UHPLC coupled with a QTRAP 5500 LC MS/MS (AB Sciex), and an Agilent 7890B/5977B GC/MS were used for metabolite analysis. The data were analyzed by MultiQuant 3.0.2 (AB Sciex) and Agilent MassHunter Quantitative Analysis Software^39^.

### Statistical analysis

Multivariate analysis was performed with a supervised classification model partial least-squares discriminant analysis (PLS-DA) after pareto scaling using MetaboAnalyst 5.0 (https://www.metaboanalyst.ca/). The comparison of specific metabolites was analyzed with Volcano plot with P<0.05 and fold changes >1.3 or < -1.3 (P>1.3).

## Results

### Sex differences in retinal metabolism

To study the impact of sex differences on retinal metabolites, we analyzed abundance of metabolites from mouse retinas in both fed and fasted states. Under fed conditions, a multivariate analysis with PLSDA showed a clear separation between the male and female retinas (**Fig 1A**), demonstrating sex differences in retinal metabolism. Volcano plots showed 32 metabolites increased, and 3 metabolites decreased in female retinas compared to male retinas (**Fig 1B, Supplemental Table S2**). The female retinas had fewer long-chain fatty acids (palmitate and stearic acid) and cysteine, but more increased metabolites in the metabolism of amino acid, nucleotide, and NAD(P) (**Fig 1B-C, Supplemental Table S2**). Pantothenic acid (a vitamin precursor for CoA synthesis), trigonelline (methylated nicotinic acid in NAD metabolism), aminoadipic acid (an intermediate in lysine metabolism), oxidized glutathione, NADP, NADPH, CDP and GDP were among the top increased metabolites, indicating that the female retina has more active CoA synthesis and NAD(P) metabolism.

**Figure 1.**
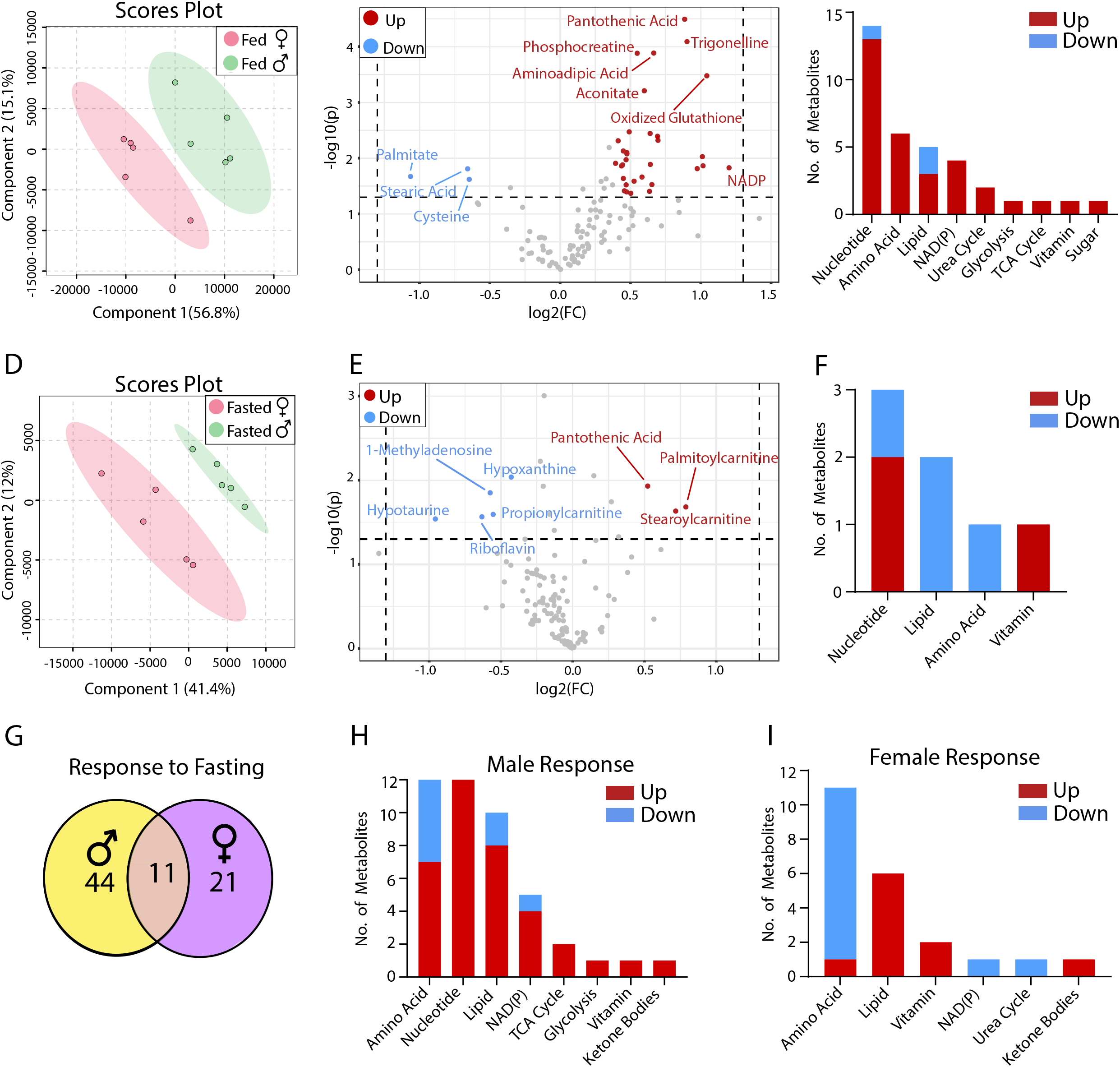
Sex difference in retinal metabolites in fed and fasted state. (A) PLSDA plots of mouse retinal metabolites from the fed state. (B) Volcano plot of retinal metabolites in the fed state. N=5. (C) The number of changed metabolites in metabolic pathways in the fed state. (E) PLSDA plots of mouse retinal metabolites from the fasted state. (F) Volcano plot of retinal metabolites in the fasted state. N=5. The number of changed metabolites from the Volcano plot in metabolic pathways in the fasted state. The number of common and sex-specific changes in retinal metabolites in response to fasting in fasted vs. fed in males or females respectively. (I) The number of changed retinal metabolites in male mice in response to fasting (J) The number of changed retinal metabolites in female mice in response to fasting.

Retinal metabolites were further separated between sexes in the PLSDA plots in the fasted state (**Fig 1D**) and sex-different metabolites were reduced to 8 when compared with fed state (**Fig1 E**). Similar to the fed state, pantothenic acid was increased in the fasted female retina. The long-chain acyl-carnitines (palmitoylcarnitine and Stearoylcarnitine) were increased, but the short-chain acyl-carnitine (propionylcarnitine), purine nucleoside (hypoxanthine and 1-methyladenosine), hypotaurine and riboflavin were decreased in the female retina (**Fig 1E-D**). Compared to the fed, 44 metabolites in the male retina and 21 in the female retina were changed in the fasted state with 11 overlapping changes between sexes (**Fig 1G, Supplemental Fig 1 and Supplemental Table S3-4**). Ketone bodies are known to increase as alternative fuels during fasting. Consistently, 3-hydroxybutyrate (3-HB) was increased 5-6 fold in both sexes after fasting. Pantothenic acid and acyl-carnitines were also increased in both sexes, suggesting that fatty acid oxidation is activated. Serine, methionine and trigonelline were decreased in the retinas of both sexes (**Fig H-I, Supplemental Table S3-4**). Despite these common changes, male and female retinas responded differently to fasting in nucleotide metabolism, amino acid metabolism, NAD(P) metabolism, TCA cycle and glycolysis (**Fig H-I, Supplemental Table S3-4**).

#### Sex difference in RPE/choroid metabolism

Similar to the neural retina, metabolites from RPE/choroid showed distinct separation between males and females in PLSDA scores plot under either fed or fasted conditions (**Fig 2-A, 2D**), indicating sex difference in RPE metabolism. The volcano plot showed 19 metabolites were significantly different between sexes under fed state, but the number of different metabolites was reduced to 9 in the fasted state (**Fig 2B, 2E, Supplemental Table S5**). Pantothenic acid was the only metabolite that was increased in the female RPE/choroid in both fed and fasted states. Succinate, 4-hydroxyproline, β-alanine and ATP were decreased in both fed and fasted states in the female (**Fig 2B, 2E**). Under the fed state, the sex-different metabolites were mostly in the metabolism of amino acid, nucleotide, TCA cycle, lipid and pentose phosphate pathway (PPP); however, under the fasted state, there were less or no changes in those pathways (**Fig 2C, 2F, Supplemental Table S5**). In response to fasting, male and female RPE/choroid showed the same number of changed metabolites with about half of them overlapping (**Fig 2G, Supplemental Fig S2, Supplemental Table S6-7**). Like the retinas, ketone bodies and acyl-carnitines were increased, whereas trigonelline and serine were decreased in both sexes in the fasted state (**Fig 2H-I, Supplemental Table S6-7**). However, female RPE/choroid had more significant changes in TCA cycle metabolites and pantothenic acid than male RPE/choroid.

**Figure 2.**
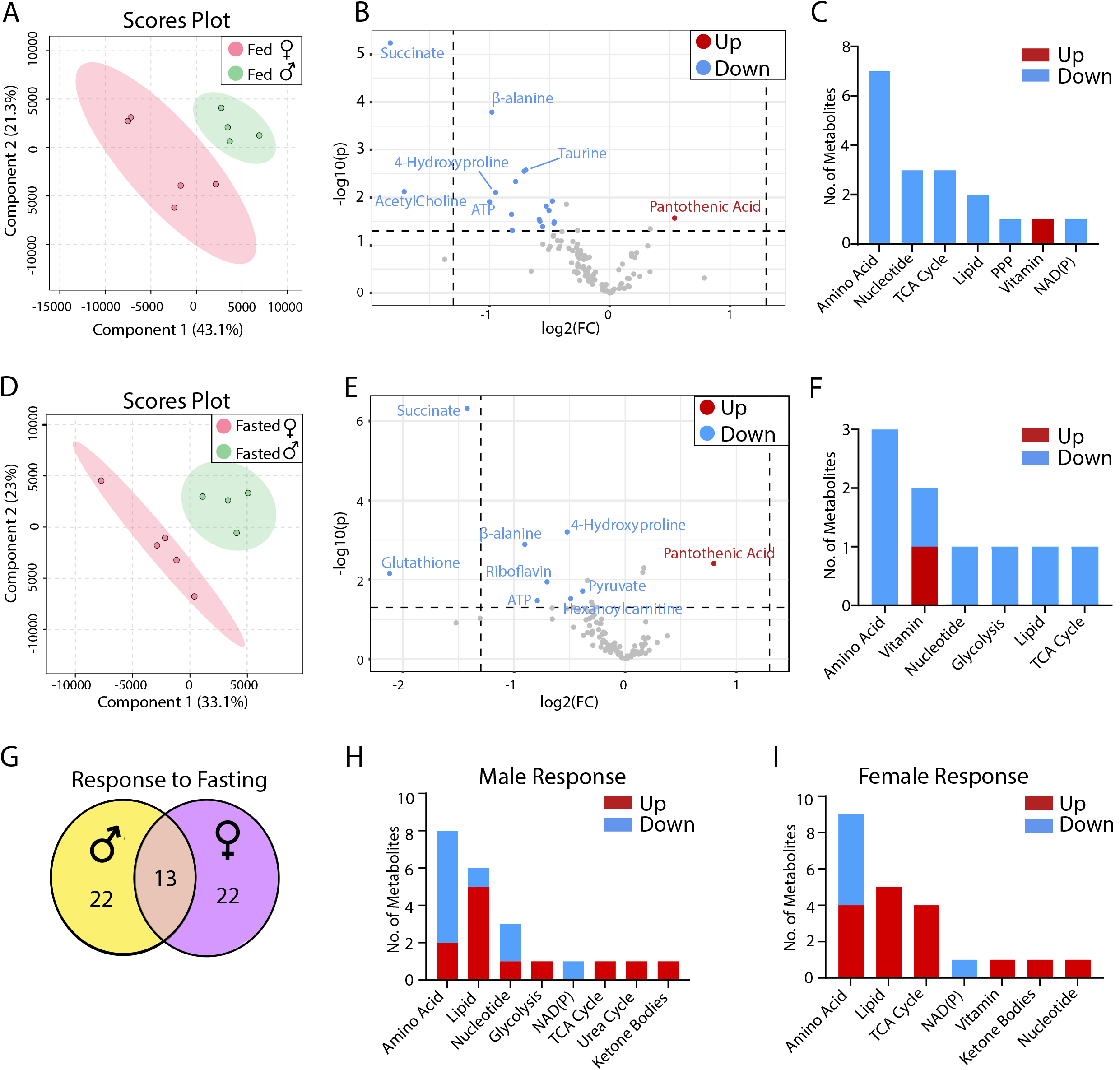
Sex difference in RPE metabolites in fed and fasted state. (A) PLSDA plots of mouse RPE metabolites from the fed state. (B) Volcano plot of RPE metabolites in the fed state. N=5. (C) The number of changed metabolites in metabolic pathways in the fed state. (E) PLSDA plots of mouse retinal metabolites from the fasted state. (F) Volcano plot of retinal metabolites in the fasted state. N=5. (G) The number of changed metabolites from the Volcano plot in metabolic pathways in the fasted state. (H) The number of common and sex-specific changes in RPE metabolites in response to fasting in fasted vs. fed in males or females respectively. (I) The number of changed RPE metabolites in male mice in response to fasting (J) The number of changed RPE metabolites in female mice in response to fasting.

#### Sex difference in Lens metabolism

PLSDA plot showed slight overlapping under the fed state but a clear separation of male and female lens metabolites in the fasted state (**Fig 3A, 3D**). Forty-five metabolites were different in the fed state and 28 under the fasted (**Fig 3B, 3E, Supplemental Table S8-9**). Twenty-two metabolites were different between males and females, independent of metabolic states. The female lens had higher glucose but lower anti-oxidative metabolites, including cystine, glutathione and ascorbic acid, than the male, suggesting female lens may be more vulnerable to oxidative stress (**Fig 3B, 3E, Supplemental Table S8-9**). Fasting-induced changes of 20 metabolites in the male and 19 in the female, with 12 metabolites changed in both sexes (**Fig 3G, Supplemental Fig 3, Supplemental Table 10-11**). Like retina and RPE, fasting increased 3-HB but decreased trigonelline and serine in the lens in both sexes. In the fasted lens, changed metabolites were primarily enriched in amino acid metabolism in both sexes. The male lens had more changes in nucleotide metabolism, while the female lens had more changes in lipid metabolism (**Fig 3H-I, Supplemental Table S10-11**).

**Figure 3.**
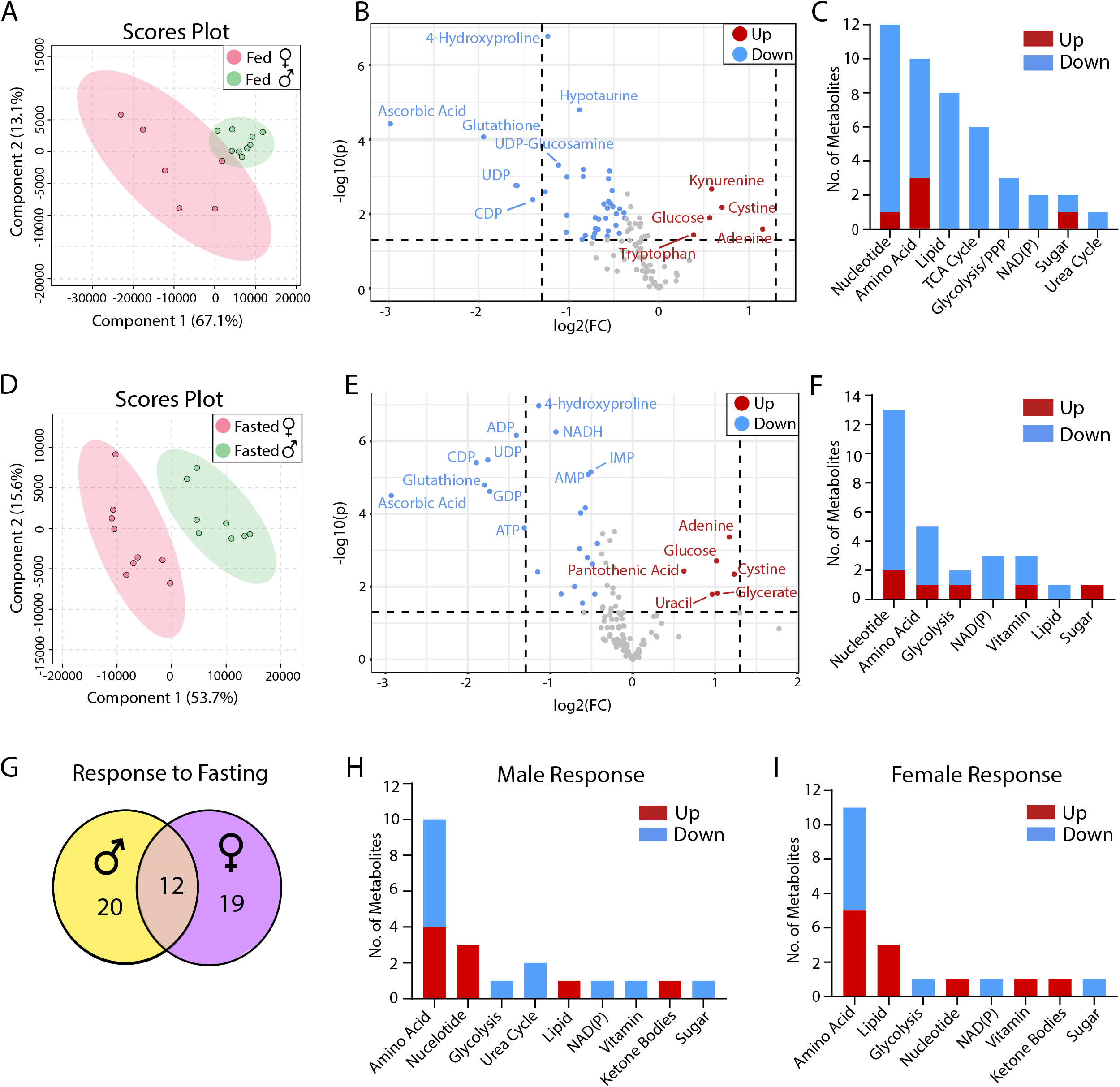
Sex difference in lens metabolites in fed and fasted state. (A) PLSDA plots of mouse lens metabolites from the fed state. (B) Volcano plot of lens metabolites in the fed state. N>6. (C) The number of changed metabolites in metabolic pathways in the fed state. (E) PLSDA plots of mouse lens metabolites from the fasted state. (F) Volcano plot of lens metabolites in the fasted state. N=9. (G) The number of changed metabolites from the Volcano plot in metabolic pathways in the fasted state. (H) The number of common and sex-specific changes in lens metabolites in response to fasting in fasted vs. fed in males or females respectively. (I) The number of changed lens metabolites in male mice in response to fasting (J) The number of changed lens metabolites in female mice in response to fasting.

#### Sex difference in brain metabolism

To reduce variation from different brain regions, we homogenized the whole brain to measure metabolites from the aliquot. PLSDA scores plot showed distinct separations of metabolites from male and female brains under either fed or fasted states (**Fig 4A, 4D**). Fourteen sex-different metabolites were in the fed state and 15 in the fasted state (**Fig 4B, 4E, Supplemental Table S12-13**). Pantothenic acid was higher and hypotaurine was lower in the female brain regardless of metabolic states. Strikingly, the sex-different metabolites were highly enriched in glucose metabolism including glycolysis and glycogen, in the fed state but not in the fasted state (**Fig 4C, 4F)**. The female brain was more sensitive to fasting than the male brain and had three times more changes in metabolites after fasting (**Fig 4G, Supplemental Fig 4, Supplemental Table S14-15**). Similar to eye tissues, 3-HB was increased and trigonelline decreased in both sexes in fasted brains. However, the female brain had massive changes of metabolites in amino acid and glucose metabolism but not the male brain (**Fig 4H-I, Supplemental Table 14-15**). These results suggest that the female brain is more sensitive to fasting and more flexible in fuel utilization than the male brain.

**Figure 4.**
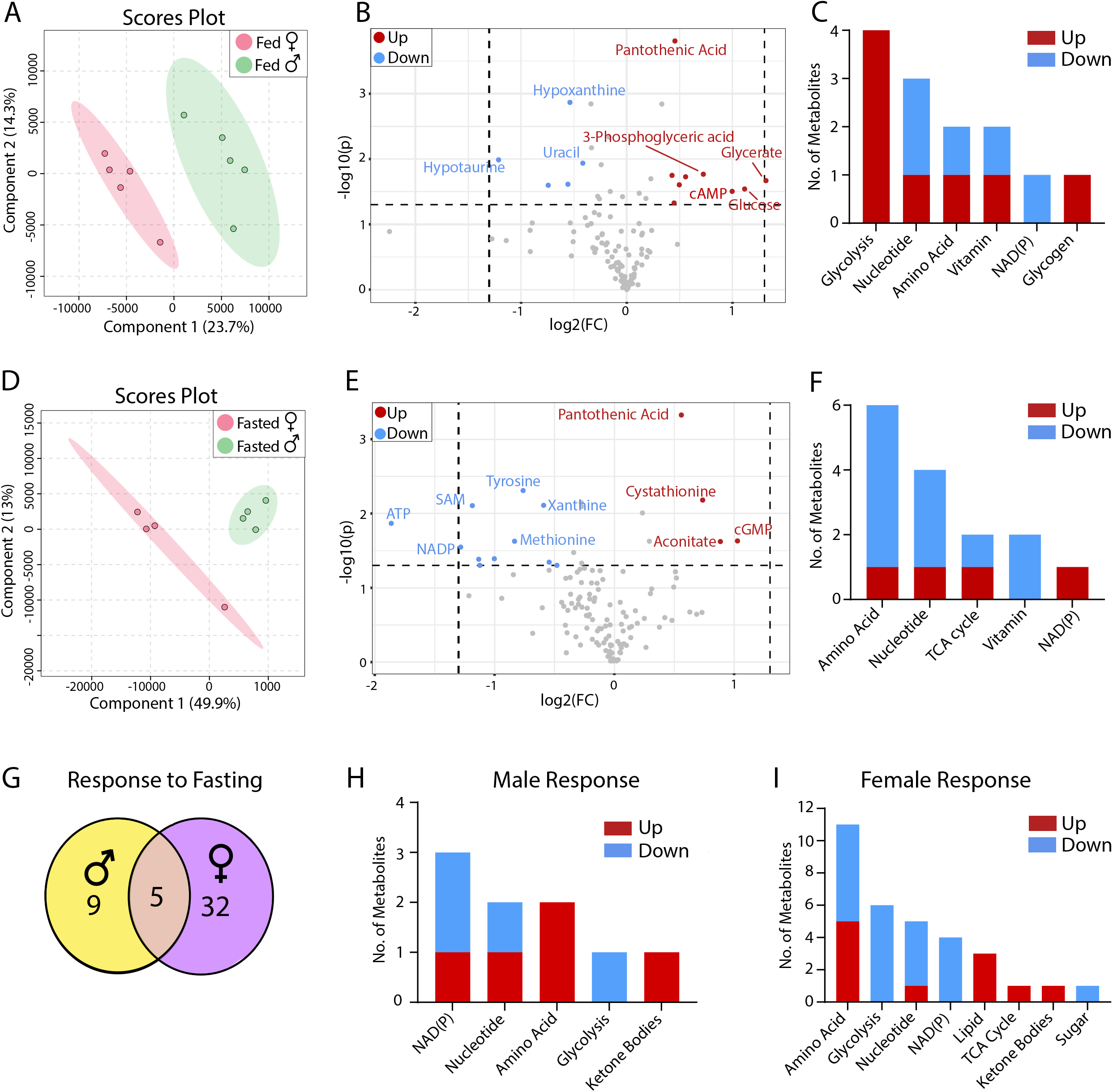
Sex difference in brain metabolites in fed and fasted state. (A) PLSDA plots of mouse brain metabolites from the fed state. (B) Volcano plot of retinal metabolites in the fed state. N=5. (C) The number of changed metabolites in metabolic pathways in the fed state. (E) PLSDA plots of mouse brain metabolites from the fasted state. (F) Volcano plot of brain metabolites in the fasted state. N=4. (G) The number of changed metabolites from the Volcano plot in metabolic pathways in the fasted state. (H) The number of common and sex-specific changes in retinal metabolites in response to fasting in fasted vs. fed males or females, respectively. (I) The number of changed retinal metabolites in male mice in response to fasting (J) The number of changed retinal metabolites in female mice in response to fasting.

#### Sex difference in plasma metabolites

We analyzed plasma metabolites to investigate whether the sex-different metabolites in the eye and brain are from the circulation. Scores plot showed minor overlapping between male and female plasma in the fed state but clear separation in the fasted state (**Fig 5A, 5D**). Sex-different metabolites from plasma were much less than those from the eye and brain, with 9 and 7 sex-different metabolites in the fed and fasted states, respectively (**Fig 5B, 5E**). Like brain and eye tissues, pantothenic acid was higher in the female plasma in the fed state, suggesting that pantothenic acid is a common sex-different metabolite. 4-hydroxyproline, pyroglutamic acid and aconitic acid were lower in the female than male plasma in either fed or fasting states (**Fig 5B, 5E**). Sex-different plasma metabolites were enriched in amino acid metabolism in both states, but the female plasma had overall more decreased metabolites than the male, especially in the fasted state (**Fig 5C, 5F**). In response to fasting, female plasma had slightly more changed metabolites than males, with less than half of overlapping changes (**Fig 5G**). Plasma glucose was significantly decreased after fasting in the data from both glucometer and targeted metabolomics.

**Figure 5.**
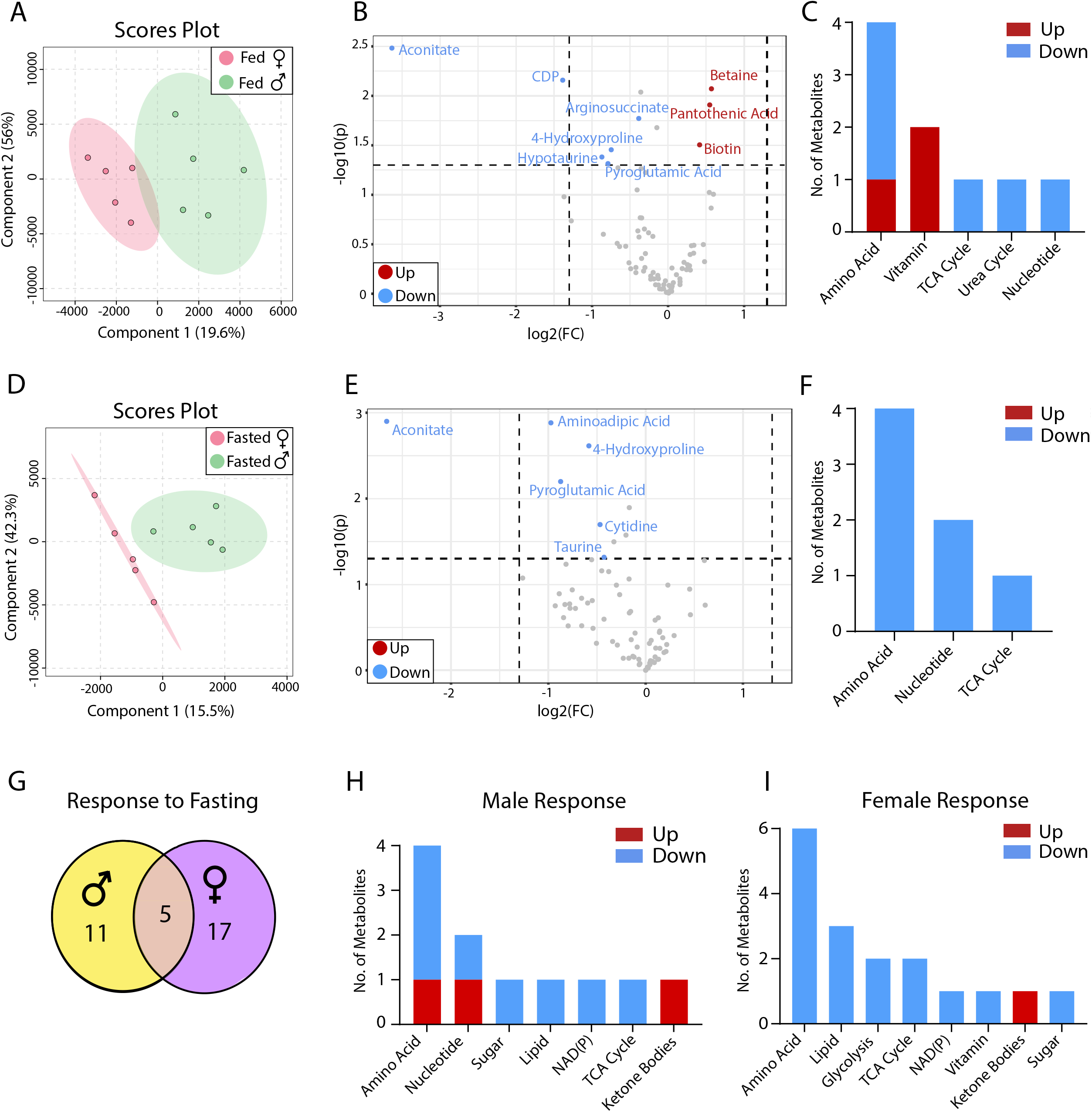
Sex difference in plasma metabolites in fed and fasted state. (A) PLSDA plots of mouse plasma metabolites from the fed state. (B) Volcano plot of plasma metabolites in the fed state. N=5. (C) The number of changed metabolites in metabolic pathways in the fed state. (E) PLSDA plots of mouse plasma metabolites from the fasted state. (F) Volcano plot of plasma metabolites in the fasted state. N=5. The number of changed metabolites from the Volcano plot in metabolic pathways in the fasted state. The number of common and sex-specific changes in plasma metabolites in response to fasting in fasted vs. fed males or females, respectively. (I) The number of changed retinal metabolites in male mice in response to fasting (J) The number of changed plasma metabolites in female mice in response to fasting.

However, there was no difference in males and females in either metabolic state (**Supplemental 5, Supplemental Table S16-17**). 3-HB and trigonelline changed similarly to other tissues after fasting, demonstrating that these two metabolites are common fasting-sensitive metabolites (**Fig 5H-I, Supplemental Fig 6 and Supplemental Table S16-17**). Both sexes were enriched in the changes of amino acid metabolism, but the female had more decreased metabolites after fasting (**Fig 5H-I**). Overall, these results suggest that except for several metabolites such as pantothenic acid, 3-HB and trigonelline, most of the sex-specific metabolic changes in the tissues are not from the systemic circulation.

#### Common and tissue-specific metabolic changes in different sexes

Thirty-two metabolites were commonly changed in two or more tissues and 64 metabolites were tissue-specific (**Fig 6A-B**). There were more common and tissue-specific metabolite changes in the fed than in fasted states. Pantothenic acid, hypotaurine and 4-hydroxyproline were the top commonly changed metabolites. Under the fed state, the lens and retina had the most tissue-specific sex-different metabolites, followed by RPE, brain and blood (**Fig 6B**). Under the fasted state, the number of these tissue-specific sex-different metabolites was reduced 2-9-fold in the lens, RPE and retina and lens whereas the tissue-specific metabolites were increased in the brain and plasma, demonstrating that the response to fasting is sex-specific and tissue-specific.

**Figure 6.**
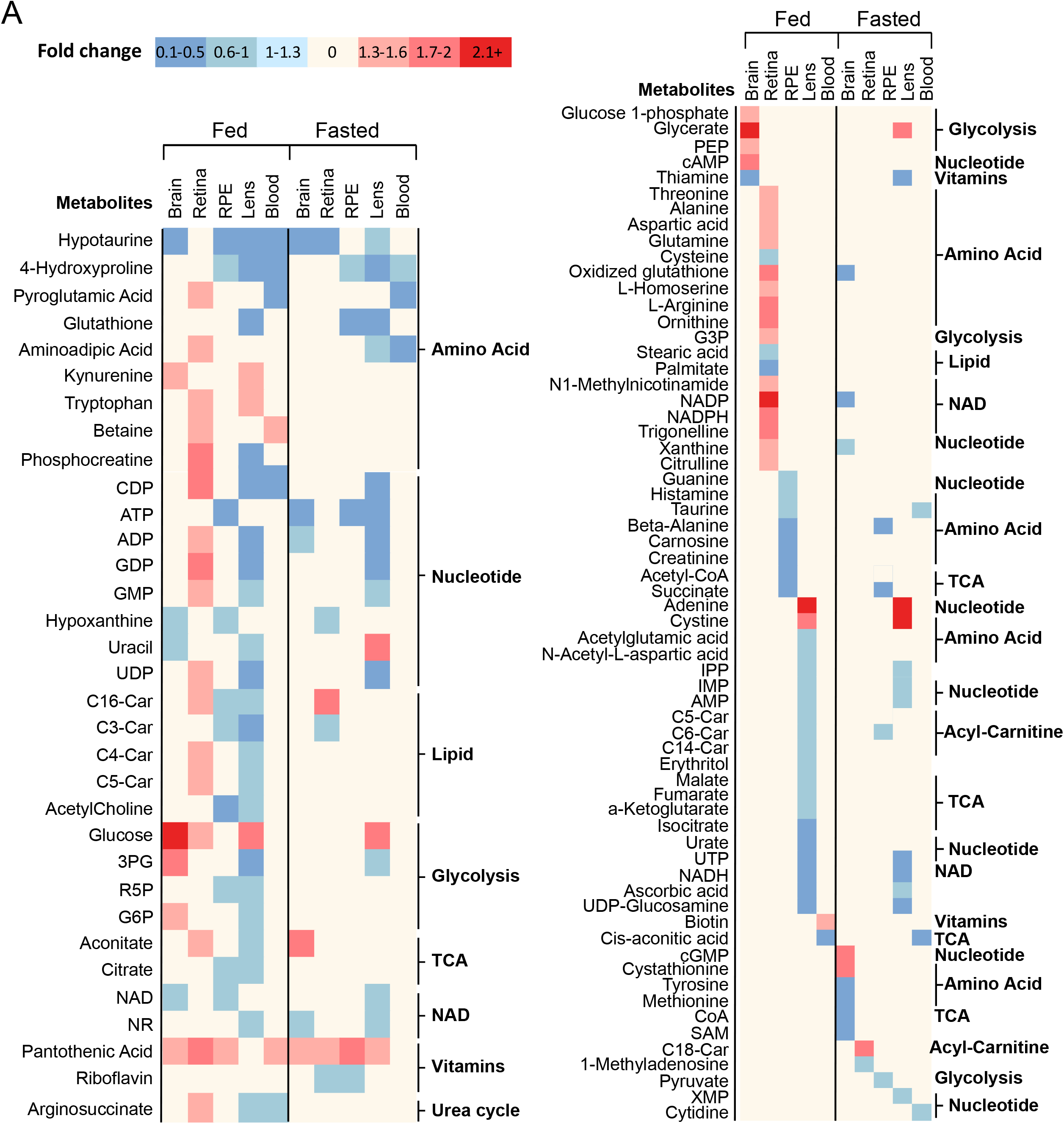
Common and tissue-specific changes of metabolites in different sexes. (A) A heat map of sex-different metabolites in two tissues and more in either fed or fasted state. (B) A heat map of tissue-specific sex difference in metabolites in fed or fasted. C3-Car, proliponylcarnitine; C4-Car, Butyrylcarnitine; C5-Car, 2-Methylbutyroylcarnitine; C6-Car, Hexanoylcarnitine; C14-Car, Myristoylcarnitine; C16-Car, Palmitoylcarnitine; C18-Car, Stearoylcarnitine; 3PG, 3-Phosphoglyceric acid; R5P, Ribose 5-phosphate; G6P, Glucose 6-phosphate; PEP, 2-phosphoenolpyruvate; G3P, Glycerol-3-phosphate; IPP, Isopentenyl pyrophosphate; SAM, S-adenosylmethionine; XMP, Xanthosine monophosphate.

To reveal common and tissue and sex-specific responses to fasting, we grouped changed metabolites of different tissues after fasting in either male or female into commonly changed (>2 tissues in either sex, 62 metabolites) and tissue-specific changed metabolites (40 metabolites) (**Fig 7A-B**). After fasting, 3-HB was increased, but trigonelline was reduced in all the tissues. Fasting reduced the level of glucose in blood, the lens and the female brain but not the retina and RPE. However, the retina and RPE had more pronounced changes of acyl-carnitines than other tissues (**Fig 7A**). The male retina and female brain had the highest number of tissue-specific metabolites, with 13 and 9, respectively. However, the female retina and male brain only had 2 and 1 tissue-specific changes (**Fig 7B**), further supporting that there is a robust tissue-specific sex difference in metabolic response to fasting.

**Figure 7.**
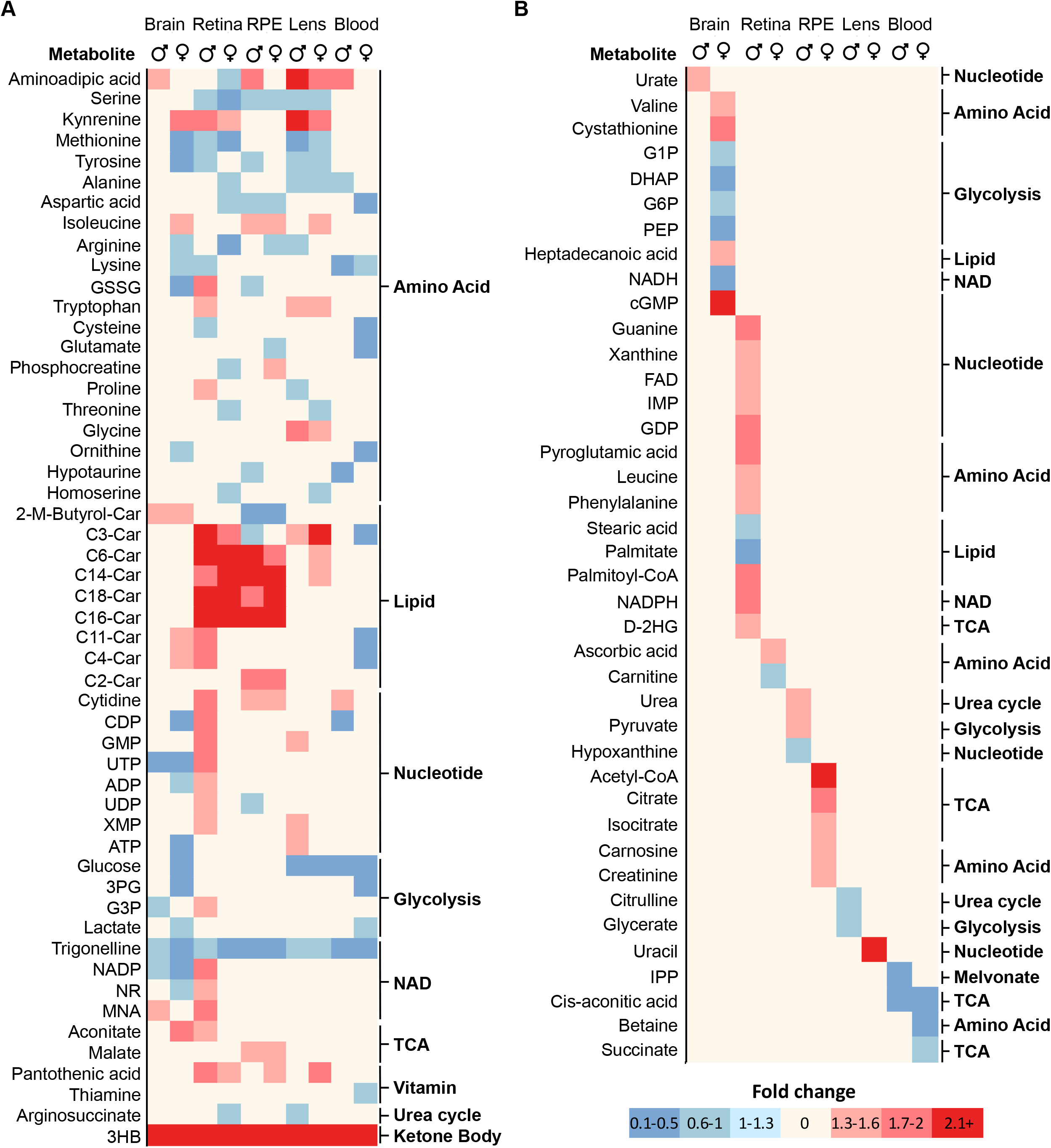
Common and sex-specific metabolites in different tissues in response to fasting. (A) A heat map of changed metabolites in two tissues and more in response to fasting in either males or females. (B) A heat map of tissue-specific changes of metabolites in response to fasting in either males or females. GSSG, Oxidized Glutathione; C11-Car, 4,8-dimethylnonanoylcarnitine; C2-Car, L-acetylcarnitine; NMA, N1-Methylnicotinamide; 3-HB, 3-hydroxybutyrate; G1P, Glucose-1-Phosphate; DHAP, Dihydroxyacetone phosphate; G6P, Glucose-6-Phosphate; D-2HG, D-2-Hydroxyglutarate.

## DISCUSSION

In this study, we have found common and tissue-specific sex differences in the metabolome of the eye and brain under different metabolic states. Pantothenic acid, a primary precursor of CoA, is a common female-enriched metabolite. Brain shows more sex-different metabolites in glycolysis, while ocular tissues show more differences in the metabolism of amino acid, lipid, nucleotide and TCA cycle. We also found male retina, female brain and female RPE are more sensitive to nutrient deprivation. Our results suggest that there are fundamental sex differences in eye metabolism.

### Sex-different metabolites in both eye and brain

Neurodegenerative diseases including Alzheimer’s disease, Parkinson’s disease, multiple sclerosis and motor neuron disease, often show sexual dimorphisms^40, 41^. Interestingly, many of these neurodegenerative disorders in the brain manifest earlier morphological or pathological changes in the eye, suggesting an intrinsic connection between the eye and brain^42, 43^. Our study also showed that eye tissues and the brain shared 13 sex-different metabolites. Pantothenic acid is higher in the female brain, eye and plasma, suggesting it is a common sex-different metabolite. In a human adult metabolomics study, pantothenic acid is increased in the female urine^44^.Interestingly, pantothenic acid and CoA-dependent mitochondrial enzymes are decreased in brain regions of patients with Alzheimer’s disease^45, 46^. However, dietary pantothenic acid intake is associated with increased cerebral amyloid β burden in patients with cognitive impairment^47^. It will be interesting to investigate the role of pantothenic acid in sexual dimorphisms in neurological diseases and their ocular symptoms.

Remarkably, the lens shows more overlapping changes with the brain, particularly in glucose metabolism, than with the neural retina and RPE. Recent studies showed extensive similarities between neurons and lens fiber cells in cell morphology and gene expression^48, 49^. Similar to the brain, lens metabolism highly depends on glucose and the deficiency of glucose transporter 1 in lens epithelium can lead to cataract formation^50^

### Tissue-specific, Sex-different metabolites

Sex hormones play critical roles in regulating energy metabolism by modulating substrate metabolism, the permeability of retina- and brain-blood barrier, transcriptional factor bindings and epigenetic regulation^34, 40, 51-53^. The different signaling of sex hormones in the retina, RPE, lens and brain regions^52, 54, 55^ may lead to differential gene expressions and nutrient availability. The differential expression of genes from sex chromosomes, sex-different sensitivity to insulin and different levels of adipokines can also contribute to tissue-specific metabolism in different metabolic states^56-59^.

The retina is metabolically demanding to maintain active visual transduction and renew daily shed outer segments^23, 60^. Retina primarily uses glucose but also prefers glutamate and aspartate for its metabolism^22, 36^. Female retinas have higher availability of amino acids including aspartate, short-chain acyl-carnitines, and metabolites in NAD(P)(H) metabolism, probably accounting for much fewer metabolite changes upon fasting compared to the male retinas. It remains to be determined the mechanisms that cause the higher nutrient availability in the female retina and their implications in the sex difference in retinal physiology and diseases.

Unlike the neural retina, RPE shows lower levels of metabolites in the female under both fed and fasted states. These decreased metabolites are mainly amino acids (histamine, taurine, carnosine, creatinine and beta-alanine) and mitochondrial intermediates (acetyl-CoA, succinate). Taurine, carnosine and its precursor beta-alanine have beneficial anti-oxidant properties^61, 62^. The decrease of these amino acids may predispose the female RPE to oxidative damage. Consistently, Female mice show more severe RPE damage and reduced retinal thickness under oxidative damage induced by sodium iodate^63, 64^. RPE mitochondria prefer to oxidize succinate from the retina to produce malate and fumarate for the retina^65-67^. However, succinate is lower in female RPE under both fed and fasted states. The female RPE may import less succinate from the retina and circulation or oxidize more succinate, resulting in sex differences in RPE mitochondrial metabolism.

Lens has the largest number of sex-different metabolites among eye tissues. Except for cystine and adenine, most the changed metabolites including mitochondrial intermediates, purine metabolites, acyl-carnitines and amino acids, were lower in the female lens. Mouse lens transcriptome shows the sex-different gene expression in mitochondrial metabolism, amino acid transport and acyl-CoA metabolism^68^. Lens mitochondria exist only in anterior epithelial cells, providing ∼30% of energy for the entire lens^69^.

The lens epithelial cells are critical to maintaining lens transparency through nutrient transport, metabolism and synthesis. The dysfunction of lens epithelial cells can lead to female-prevalent cataracts^10^. In human donor eyes, the female has higher epithelial cell density than the male^6^. The lens metabolism relies on nutrients from the aqueous humor. There are significant sex differences in human aqueous humor proteome and protein^70, 71^. However, sex-different metabolites in aqueous humor are still unclear as most aqueous humor metabolomics are sex-matched without analysis of the sex differences ^72, 73^. Further studies on the metabolites of lens epithelial cells and aqueous humor will help understand the sex difference in lens metabolism and its implications in cataracts.

Our results show the female brain has more sensitive glucose metabolism in different metabolic states, which may implicate in the sex differences in brain metabolism, physiology and diseases. The brain relies on glucose as its primary source of energy. Human studies with positron emission tomography (PET) show that young women have high cerebral blood flow and glycolysis than men^74, 75^. Aerobic glycolysis is positively correlated to brain aging^76^ aging and adult female brain show a few years of metabolic youthfulness than the male^74^. However, this metabolic youthfulness starts to disappear in cognitively impaired patients such as Alzheimer’s disease, probably due to a higher rate of decline in glucose metabolism in the female patients^77^. Both human studies and animal models also show that one of the earliest signs of Alzheimer’s disease is a reduction of cerebral glucose metabolism and the disturbed glucose metabolism is associated with disease progression^78^. These findings are consistent with our results that female brain has higher glycolytic metabolites in the fed state and more sensitive in changes of glucose metabolism to nutritional stress, suggesting that early intervention to glucose metabolism may be important in the female under stressed conditions such as neurological diseases.

In conclusion, our findings demonstrate substantial sex differences in metabolic profiles in different eye tissues, supporting the urgent need to include animals and humans of both sexes in eye metabolomics research. We also find similarities in sex-different metabolites between the eye and brain, supporting that eye metabolomics can serve as an important window to reflect brain metabolism and diseases.

## Acknowledgment

This work was supported by National Institutes of Health Grant EY031324 (JD), EY032462(JD), Bright Focus Foundation M2020141(JD), the Retina Research Foundation (JD) and funds for Core facilities P20 GM103434 (WV INBRE grant), WVCTSI grant GM104942.

