## Supplemental Information for "Tissue-specific sex difference in mouse eye and brain metabolome under fed and fasted states"

This file includes:

Supplementary Figure S1-6

Supplementary Table S1-17

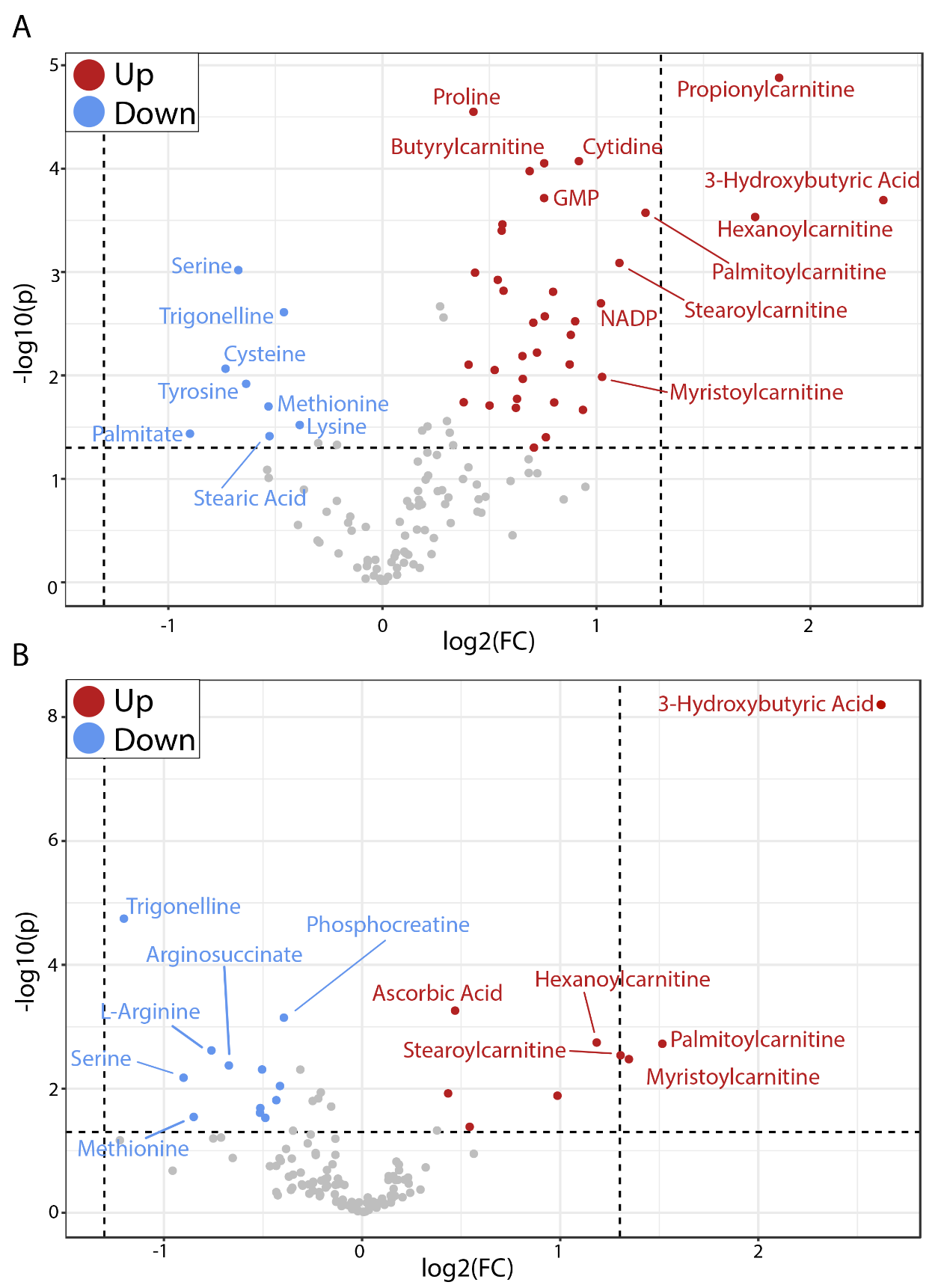

**Supplemental Figure S1. (A)** Volcano plots of retinal metabolites from fed vs. fasted male mice. **(B)** Volcano plots of retinal metabolites from fed vs. fasted female mice.

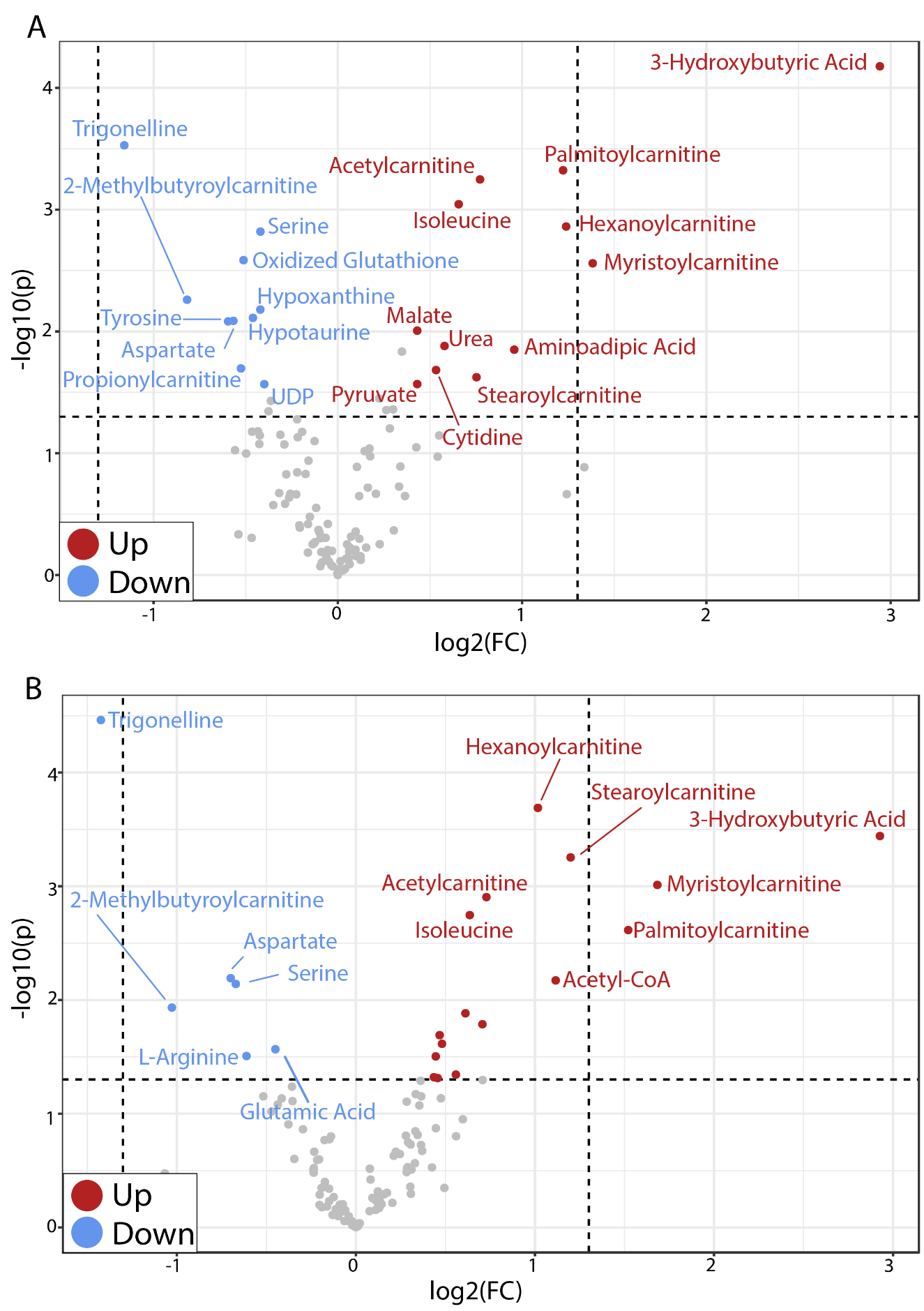
**Supplemental Figure S2. (A)** Volcano plots of RPE metabolites from fed vs. fasted male mice. **(B)** Volcano plots of RPE metabolites from fed and fasted male mice.

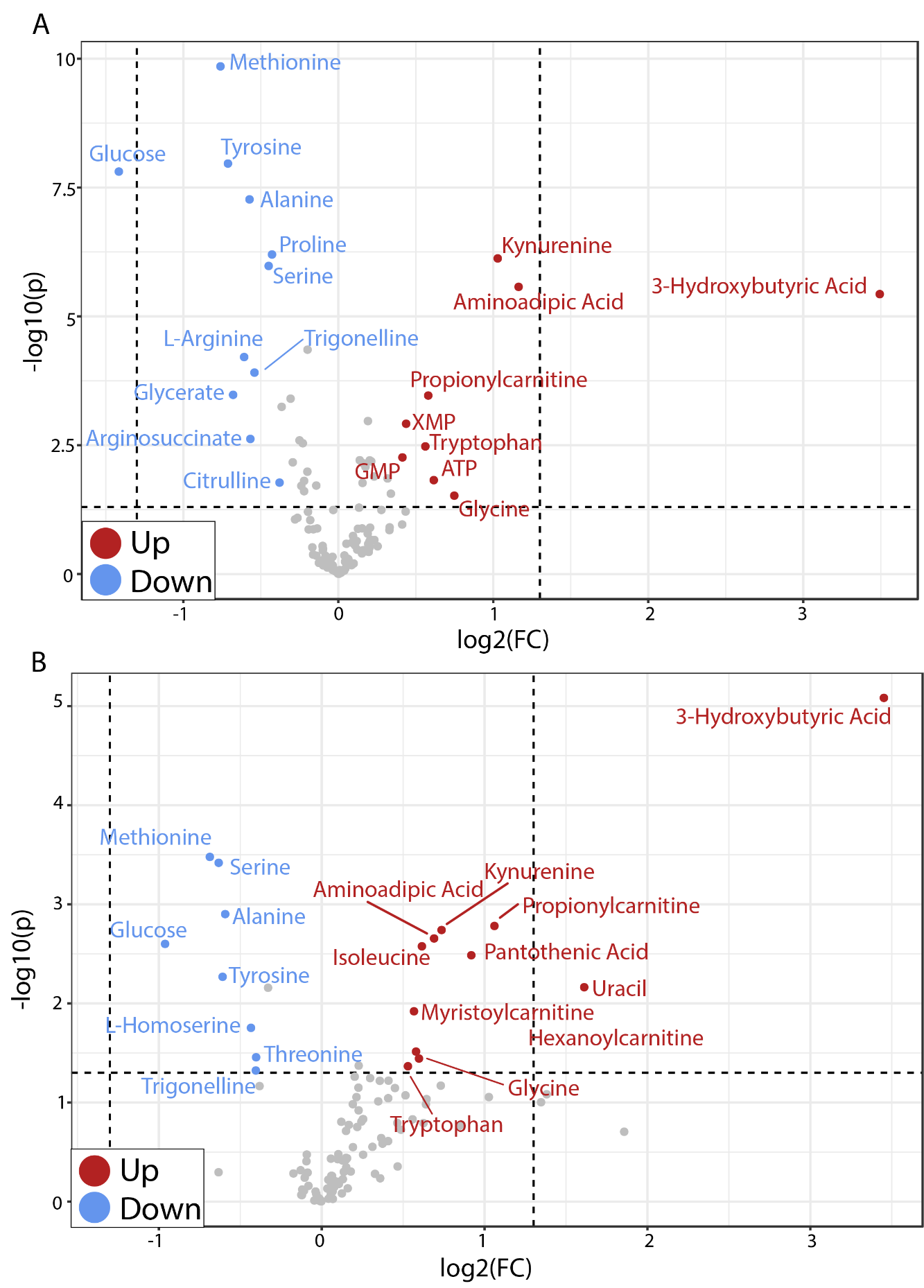

**Supplemental Figure S3. (A)** Volcano plots of lens metabolites from fed vs. fasted male mice. **(B)** Volcano plots of lens metabolites from fed and fasted male mice.

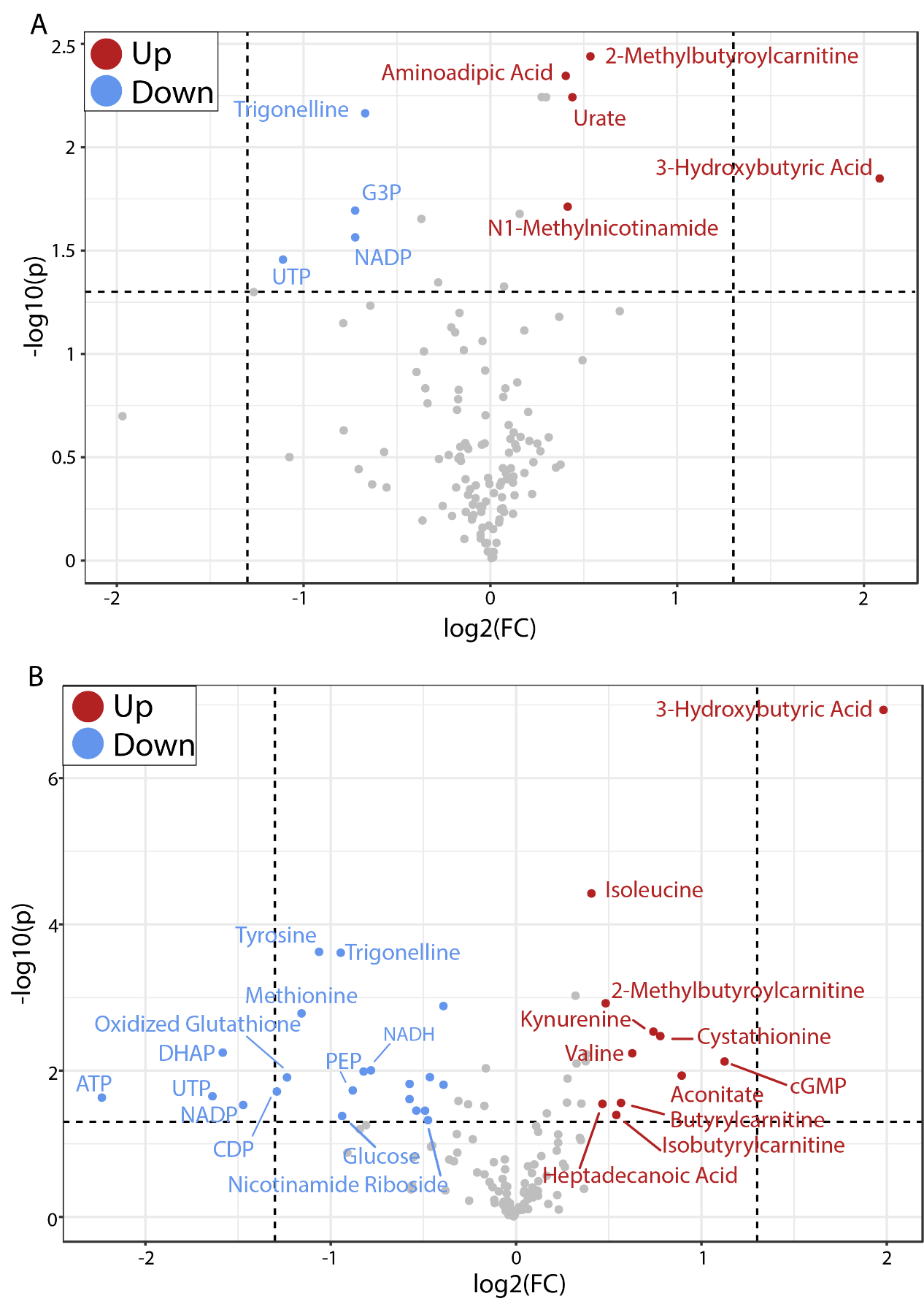

**Supplemental Figure S4. (A)** Volcano plots of brain metabolites from fed vs. fasted male mice. **(B)** Volcano plots of brain metabolites from fed and fasted male mice.

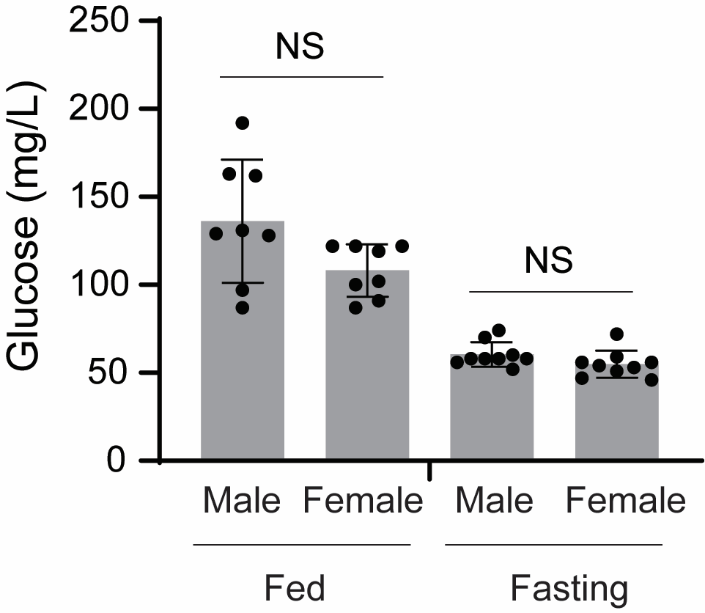

**Supplemental Figure S5.** Plasma glucose levels in fed and fasted mice. N=5 *P<0.05**.** Glucose levels were measured via tail bleeding using a glucometer (Contour blood glucose meter 9545C).

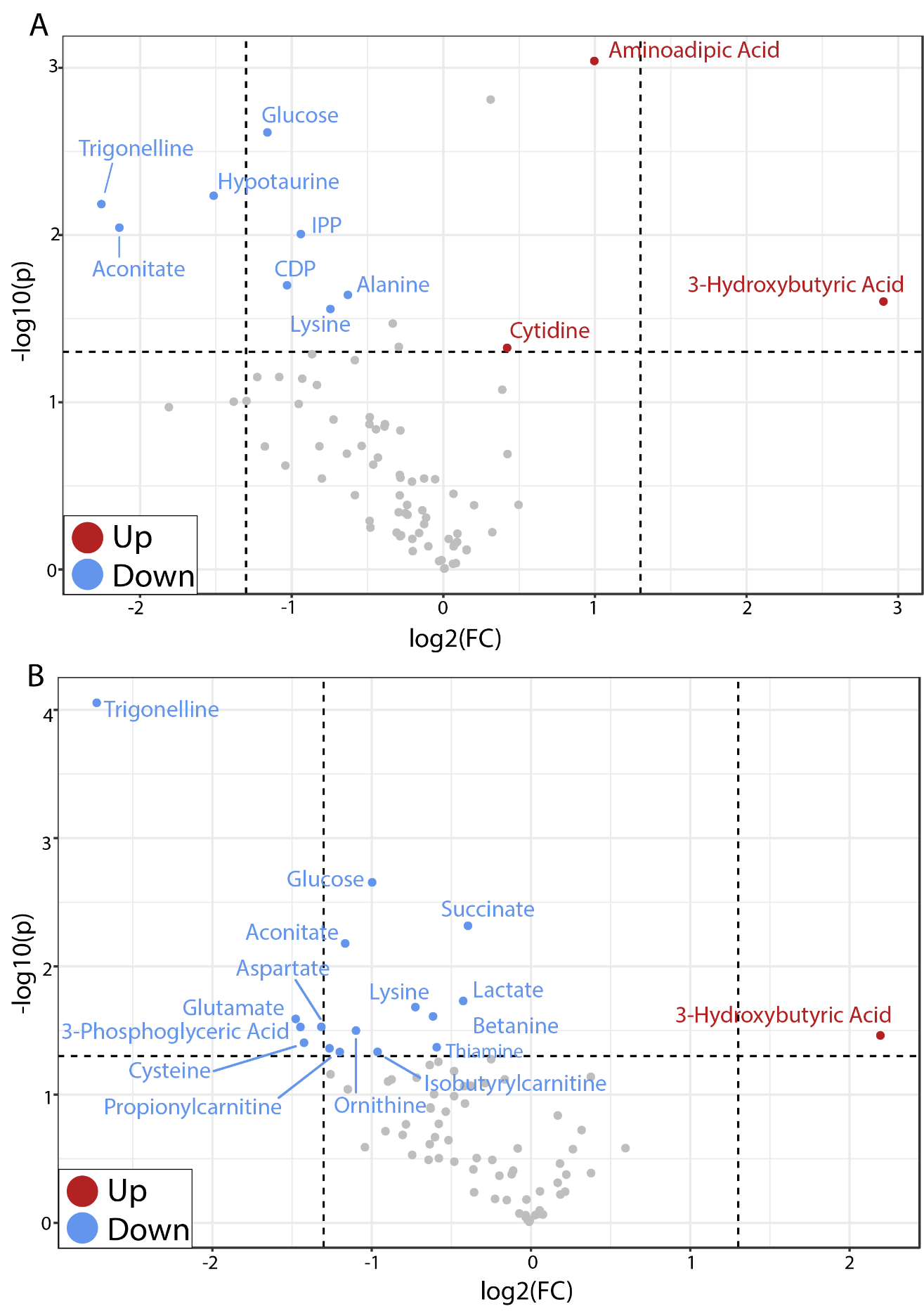

**Supplemental Figure S6 (A)** Volcano plots of plasma metabolites from fed vs. fasted male mice. **(B)** Volcano plots of plasma metabolites from fed and fasted male mice.

**Supplemental Table S1**. The lists of measured metabolites and their parameters in mass spectrometry.

| **Metabolite** | **Platform** | **Pathway** | **CAS ID** |
| --- | --- | --- | --- |
| 3-Hydroxybutyric acid | GCMS | Lipid/Ketone bodies | 300-85-6 |
| 3-Phosphoglyceric acid | GCMS | Glycolysis | 80731-10-8 |
| a-Ketoglutarate | GCMS | TCA Cycle | 328-50-7 |
| Asparagene | GCMS | Amino Acid | 70-47-3 |
| Beta-Alanine | GCMS | Amino acid | 107-95-9 |
| Biotin | GCMS | Vitamins | 58-85-5 |
| Cholesterol | GCMS | Lipid | 57-88-5 |
| Citrate | GCMS | TCA Cycle | 77-92-9 |
| Cysteine | GCMS | Amino Acid | 56-89-3 |
| Cystine | GCMS | Amino Acid | 56-89-3 |
| DHAP | GCMS | Glycolysis | 57-04-5 |
| Fumarate | GCMS | TCA Cycle | 110-17-8 |
| Glycerate | GCMS | Glycolysis | 14028-62-7 |
| Isocitrate | GCMS | TCA Cycle | 1637-73-6 |
| Isoleucine | GCMS | Amino Acid | 73-32-5 |
| L-Histidine | GCMS | Amino Acid | 332-80-9 |
| Malate | GCMS | TCA Cycle | 6915-15-7 |
| Palmitate | GCMS | Lipid metabolism/Fatty Acids | 57-10-3 |
| PEP | GCMS | Glycolysis | 138-08-9 |
| Pyroglutamic acid | GCMS | Amino Acid | 98-79-3 |
| Pyruvate | GCMS | Glycolysis | 113-24-6 |
| Stearic acid | GCMS | Lipid metabolism/Fatty Acid | 57-11-4 |
| Taurine | GCMS | Amino Acid | 107-35-7 |
| Tryptophan | GCMS | Amino Acid | 73-22-3 |
| Uracil | GCMS | Nucleotide/Pyrimidine metabolism | 66-22-8 |
| Urea | GCMS | Urea Cycle | 57-13-6 |
| 1-Methyladenosine | LCMS | Nucleotide/Purine metabolism | 15763-06-1 |
| 2-Methylbutyroylcarnitine | LCMS | AcylCarnitine/Amino acid metabolism | 31023-25-3 |
| 3-Aminoisobutanoic acid | LCMS | Nucleotide/pyrimidine (thymine) | 144-90-1 |
| 4-Hydroxyproline | LCMS | Amino Acid metabolism/Proline/Arginine | 51-35-4 |
| AcetylCarnitine | LCMS | Lipid metabolism | 14992-62-2 |
| AcetylCholine | LCMS | Lipid/phospholipid, | 51-84-3 |
| Acetyl-CoA | LCMS | TCA Cycle/Fatty acid metabolism | 72-89-9 |
| Acetylglutamic acid | LCMS | Amino Acid metabolism/Proline/Arginine | 1188-37-0 |
| Aconitate | LCMS | TCA Cycle | 499-12-7 |
| Adenine | LCMS | Nucleotide/Purine metabolism | 73-24-5 |
| Adenosine | LCMS | Nucleotide/Purine metabolism | 58-61-7 |
| ADP | LCMS | Nucleotide/Purine metabolism | 58-64-0 |
| AICAR | LCMS | Nucleotide/Purine metabolism | 3031-94-5 |
| Alanine | LCMS | Amino Acid | 56-41-7 |
| Aminoadipic acid | LCMS | Amino Acid metabolism/Lysine | 542-32-5 |
| AMP | LCMS | Nucleotide | 61-19-8 |
| Arginosuccinate | LCMS | Urea Cycle | 2387-71-5 |
| Ascorbic acid | LCMS | Ascorbate and aldarate metabolism | 50-81-7 |
| Aspartic acid | LCMS | Amino Acid | 56-84-8 |
| ATP | LCMS | Nucleotide | 987-65-5 |
| Betaine | LCMS | Amino Acid/gly,ser, thr metabolism | 590-46-5 |
| Butyrylcarnitine | LCMS | Lipid metabolism/Fatty Acid metabolism | 25576-40-3 |
| cAMP | LCMS | Nucleotide/Purine metabolism | 60-92-4 |
| Carnitine | LCMS | Amino Acid metabolism/lys | 541-15-1 |
| Carnosine | LCMS | Amino Acids/Histidine | 305-84-0 |
| CDP | LCMS | Nucleotide/Pyrimidine metabolism | 63-38-7 |
| cGMP | LCMS | Nucleotide/Purine metabolism | 7665-99-8 |
| Choline | LCMS | Lipid metabolism | 62-49-7 |
| cis-Aconitic acid | LCMS | TCA Cycle | 499-12-7 |
| Citrulline | LCMS | Urea Cycle | 372-75-8 |
| CoA | LCMS | TCA Cycle/Fatty acid metabolism | 55672-92-9 |
| Creatine | LCMS | Amino Acids/Arg, gly | 57-00-1 |
| Creatinine | LCMS | Amino Acids/Arg, gly | 60-27-5 |
| Cystathionine | LCMS | Amino Acids/cys | 56-88-2 |
| Cytidine | LCMS | Nucleotide | 65-46-3 |
| D-2-hydroxyglutarate | LCMS | TCA cycle | 103404-90-6 |
| D-Ribulose 5-phosphate | LCMS | Glycolysis/PPP | 551-85-9 |
| Erythritol | LCMS | Sugar | 149-32-6 |
| FAD | LCMS | Nucleotide | 146-14-5 |
| G3P | LCMS | Glycolysis | 142-10-9 |
| Gamma-Aminobutyric acid | LCMS | Amino Acid metabolism/Ala, Glu, Asp | 56-12-2 |
| GDP | LCMS | Nucleotide/Purine metabolism | 146-91-8 |
| Glucose | LCMS | Glycolysis/sugar | 492-62-6 |
| Glucose 1-phosphate | LCMS | Glycolysis | 59-56-3 |
| Glucose 6-phosphate | LCMS | Glycolysis/PPP | 56- 73-5 |
| Glutamate | LCMS | Amino Acid | 56-86-0 |
| Glutamine | LCMS | Amino Acid | 56-85-9 |
| Glutathione | LCMS | Amino acid metabolism | 70-18-8 |
| Glycine | LCMS | Amino Acid | 56-40-6 |
| GMP | LCMS | Nucleotide/Purine metabolism | 85-32-5 |
| Guanine | LCMS | Nucleotide/Purine metabolism | 73-40-5 |
| Guanosine | LCMS | Nucleotide | 118-00-3 |
| Heptadecanoic acid | LCMS | Lipid metabolism/Fatty Acid metabolism | 506-12-7 |
| Hexanoylcarnitine | LCMS | Acylcarnitines/Fatty Acid metabolism | 22671-29-0 |
| Histamine | LCMS | Amino acid metabolism | 51-45-6 |
| Hypotaurine | LCMS | Taurine metabolism | 300-84-5 |
| Hypoxanthine | LCMS | Nucleotide/Purine metabolism | 68-94-0 |
| IMP | LCMS | Nucleotide/Purine metabolism | 131-99-7 |
| Inosine | LCMS | Nucleotide/Purine metabolism | 58-63-9 |
| IPP | LCMS | mevalonate pathway | 18687-43-9 |
| Isobutyrylcarnitine | LCMS | Lipid metabolism/Fatty Acid metabolism | 25518-49-4 |
| Kynurenine | LCMS | Amino Acid metabolism/Trp | 343-65-7 |
| Lactate | LCMS | Glycolysis | 50-21-5 |
| L-Arginine | LCMS | Amino Acid/Urea cycle | 74-79-3 |
| L-Asparagine | LCMS | Amino Acid/ AA metabolism/Ala, Glu, Asp | 70-47-3 |
| Leucine | LCMS | Amino Acid | 61-90-5 |
| L-Homoserine | LCMS | Amino Acid/Thr, Met, Asp | 1927-25-9 |
| Lysine | LCMS | Amino Acid | 56-87-1 |
| Methionine | LCMS | Amino Acid | 63-68-3 |
| myo-Inositol | LCMS | sugar | 87-89-8 |
| Myristoylcarnitine | LCMS | Lipid metabolism/Fatty Acid metabolism | 25597-07-3 |
| N1-Methylnicotinamide | LCMS | NAD Metabolism | 3106-60-3 |
| N-Acetyl-L-aspartic acid | LCMS | Amino Acid metabolism/Ala, Glu, Asp | 997-55-7 |
| NAD | LCMS | Nicotinate and nicotinamide metabolism | 53-84-9 |
| NADH | LCMS | Nicotinate and nicotinamide metabolism | 58-68-4 |
| NADP | LCMS | Nicotinate and nicotinamide metabolism | 53-59-8 |
| NADPH | LCMS | Nicotinate and nicotinamide metabolism | 2646-71-1 |
| Niacinamide | LCMS | Nicotinate and nicotinamide metabolism | 98-92-0 |
| Nicotinamide riboside | LCMS | Nicotinate and nicotinamide metabolism | 2181-04-6 |
| Ornithine | LCMS | Amino acid/Urea cyle | 70-26-8 |
| Oxalic acid | LCMS | Glyoxylate and dicarboxylate metabolism | 144-62-7 |
| Oxidized Glutathione | LCMS | Amino acid metabolism | 13081-14-6 |
| Palmitoylcarnitine | LCMS | Acylcarnitines/Fatty Acid metabolism | 1985-18-8 |
| Palmitoyl-CoA | LCMS | Lipid metabolism/Fatty Acid metabolism | 188174-64-3 |
| Pantothenic acid | LCMS | Vitamin/CoA | 137-08-6 |
| Phenylalanine | LCMS | Amino Acid | 63-91-2 |
| Phosphocreatine | LCMS | Amino Acid/pro and arginine | 19333-65-4 |
| Proline | LCMS | Amino Acid | 4298-08-2 |
| Propionylcarnitine | LCMS | Acylcarnitine/Fatty Acid metabolism | 17298-37-2 |
| Riboflavin | LCMS | Vitamins | 83-88-5 |
| SAM | LCMS | Amino Acid/pro and arginine | 86867-01-8 |
| Serine | LCMS | Amino Acid | 56-45-1 |
| Stearoylcarnitine | LCMS | Acylcarnitines/Fatty Acid metabolism | 25597-09-5 |
| Succinate | LCMS | TCA Cycle | 110-15-6 |
| Thiamine | LCMS | Vitamins | 59-43-8 |
| Threonine | LCMS | Amino Acid | 72-19-5 |
| Trigonelline | LCMS | Nicotinate and nicotinamide metabolism | 535-83-1 |
| Tyrosine | LCMS | Amino Acid | 60-18-4 |
| UDP | LCMS | Nucleotide/Pyrimidine metabolism | 58-98-0 |
| UDP-Glucosamine | LCMS | Nucleotide sugar | 17479-04-8 |
| Urate | LCMS | Nucleotide/Purine metabolism | 69-93-2 |
| Uridine diphosphate glucose | LCMS | Nucleotide sugar | 133-89-1 |
| UTP | LCMS | Nucleotide/pyrimidine | 63-39-8 |
| Valine | LCMS | Amino Acid | 72-18-4 |
| Xanthine | LCMS | Nucleotide | 69-89-6 |
| Xanthosine | LCMS | Nucleotide/Purine metabolism | 146-80-5 |
| XMP | LCMS | Nucleotide/Purine metabolism | 523-98-8 |

**Supplemental Table S2**. Sex different retinal metabolites in different metabolic pathways in the fed state by Volcano plots.

| **Metabolite** | **Pathway** | **FC** | **log2(FC)** | **P Value** | **-log10(p)** |
| --- | --- | --- | --- | --- | --- |
| Propionylcarnitine | Acylcarnitine/Fatty Acid metabolism | 3.6136 | 1.8534 | 1.32E-05 | 4.8787 |
| Proline | Amino Acid | 1.343 | 0.42551 | 2.82E-05 | 4.5498 |
| Cytidine | Nucleotide | 1.8893 | 0.91789 | 8.47E-05 | 4.0723 |
| Butyrylcarnitine | Lipid metabolism/Fatty Acid metabolism | 1.6889 | 0.75604 | 8.87E-05 | 4.0521 |
| Isobutyrylcarnitine | Lipid metabolism/Fatty Acid metabolism | 1.6111 | 0.68803 | 0.00010577 | 3.9756 |
| GMP | Nucleotide/Purine metabolism | 1.6883 | 0.75558 | 0.00019268 | 3.7152 |
| 3-Hydroxybutyric acid | Lipid/Ketone bodies | 5.0645 | 2.3404 | 0.00020165 | 3.6954 |
| Palmitoylcarnitine | Acylcarnitines/Fatty Acid metabolism | 2.3445 | 1.2293 | 0.00026731 | 3.573 |
| Hexanoylcarnitine | Acylcarnitines/Fatty Acid metabolism | 3.3436 | 1.7414 | 0.00029345 | 3.5325 |
| Leucine | Amino Acid | 1.4752 | 0.56094 | 0.00034592 | 3.461 |
| XMP | Nucleotide/Purine metabolism | 1.4717 | 0.55746 | 0.00039787 | 3.4003 |
| Stearoylcarnitine | Acylcarnitines/Fatty Acid metabolism | 2.1546 | 1.1074 | 0.00081669 | 3.0879 |
| Serine | Amino Acid | 0.6272 | -0.67296 | 0.00095775 | 3.0187 |
| Phenylalanine | Amino Acid | 1.3499 | 0.43286 | 0.0010143 | 2.9938 |
| Aconitate | TCA Cycle | 1.4528 | 0.53887 | 0.0011904 | 2.9243 |
| IMP | Nucleotide/Purine metabolism | 1.4803 | 0.56588 | 0.0015179 | 2.8187 |
| Pantothenic acid | Vitamin/CoA | 1.7378 | 0.79725 | 0.0015518 | 2.8092 |
| NADP | Nicotinate and nicotinamide metabolism | 2.0291 | 1.0209 | 0.002006 | 2.6977 |
| Trigonelline | Nicotinate and nicotinamide metabolism | 0.727 | -0.46005 | 0.0024475 | 2.6113 |
| Oxidized glutathione | Amino acid metabolism | 1.6918 | 0.75859 | 0.0026826 | 2.5714 |
| CDP | Nucleotide/Pyrimidine metabolism | 1.8668 | 0.90055 | 0.0029955 | 2.5235 |
| N1-Methylnicotinamide | NAD Metabolism | 1.6302 | 0.70508 | 0.0030881 | 2.5103 |
| NADPH | Nicotinate and nicotinamide metabolism | 1.8407 | 0.88023 | 0.0040566 | 2.3918 |
| Pyroglutamic acid | Amino Acid | 1.6497 | 0.7222 | 0.0060116 | 2.221 |
| Tryptophan | Amino Acid | 1.5738 | 0.65422 | 0.0064967 | 2.1873 |
| GDP | Nucleotide/Purine metabolism | 1.8341 | 0.87507 | 0.0078247 | 2.1065 |
| G3P | Glycolysis | 1.322 | 0.40268 | 0.0078706 | 2.104 |
| Cysteine | Amino Acid | 0.6018 | -0.73268 | 0.008608 | 2.0651 |
| UDP | Nucleotide/Pyrimidine metabolism | 1.4385 | 0.52452 | 0.0088668 | 2.0522 |
| Myristoylcarnitine | Lipid metabolism/Fatty Acid metabolism | 2.0372 | 1.0266 | 0.010333 | 1.9858 |
| Xanthine | Nucleotide | 1.5758 | 0.65605 | 0.010804 | 1.9664 |
| Tyrosine | Amino Acid | 0.6433 | -0.63648 | 0.01205 | 1.919 |
| Nicotinamide Riboside | Nicotinate and nicotinamide metabolism | 1.5466 | 0.62907 | 0.016849 | 1.7734 |
| D-2-hydroxyglutarate | TCA cycle | 1.3012 | 0.37988 | 0.018187 | 1.7402 |
| UTP | Nucleotide/pyrimidine | 1.7443 | 0.80261 | 0.018249 | 1.7388 |
| ADP | Nucleotide/Purine metabolism | 1.4149 | 0.50071 | 0.019531 | 1.7093 |
| Methionine | Amino Acid | 0.6916 | -0.53208 | 0.019937 | 1.7003 |
| FAD | Nucleotide | 1.541 | 0.62388 | 0.020585 | 1.6864 |
| Palmitoyl-CoA | Lipid metabolism/Fatty Acid metabolism | 1.9143 | 0.93683 | 0.021527 | 1.667 |
| Lysine | Amino Acid | 0.7653 | -0.38599 | 0.030136 | 1.5209 |
| Palmitate | Lipid metabolism/Fatty Acids | 0.5363 | -0.89884 | 0.036594 | 1.4366 |
| Stearic acid | Lipid metabolism/Fatty Acid | 0.6938 | -0.5274 | 0.038629 | 1.4131 |
| Kynurenine | Amino Acid metabolism/Trp | 1.6981 | 0.76389 | 0.039618 | 1.4021 |
| Guanine | Nucleotide/Purine metabolism | 1.6336 | 0.70809 | 0.049879 | 1.3021 |

**Supplemental Table S3**. Changed retinal metabolites in different metabolic pathways in fed vs. fasted male mice by Volcano plots.

| **Metabolite** | **Pathway** | **FC** | **log2(FC)** | **P Value** | **-log10(p)** |
| --- | --- | --- | --- | --- | --- |
| Propionylcarnitine | Acylcarnitine/Fatty Acid metabolism | 3.6136 | 1.8534 | 1.32E-05 | 4.8787 |
| Proline | Amino Acid | 1.343 | 0.42551 | 2.82E-05 | 4.5498 |
| Cytidine | Nucleotide | 1.8893 | 0.91789 | 8.47E-05 | 4.0723 |
| Butyrylcarnitine | Lipid metabolism/Fatty Acid metabolism | 1.6889 | 0.75604 | 8.87E-05 | 4.0521 |
| Isobutyrylcarnitine | Lipid metabolism/Fatty Acid metabolism | 1.6111 | 0.68803 | 0.00010577 | 3.9756 |
| GMP | Nucleotide/Purine metabolism | 1.6883 | 0.75558 | 0.00019268 | 3.7152 |
| 3-Hydroxybutyric acid | Lipid/Ketone bodies | 5.0645 | 2.3404 | 0.00020165 | 3.6954 |
| Palmitoylcarnitine | Acylcarnitines/Fatty Acid metabolism | 2.3445 | 1.2293 | 0.00026731 | 3.573 |
| Hexanoylcarnitine | Acylcarnitines/Fatty Acid metabolism | 3.3436 | 1.7414 | 0.00029345 | 3.5325 |
| Leucine | Amino Acid | 1.4752 | 0.56094 | 0.00034592 | 3.461 |
| XMP | Nucleotide/Purine metabolism | 1.4717 | 0.55746 | 0.00039787 | 3.4003 |
| Stearoylcarnitine | Acylcarnitines/Fatty Acid metabolism | 2.1546 | 1.1074 | 0.00081669 | 3.0879 |
| Serine | Amino Acid | 0.62722 | -0.67296 | 0.00095775 | 3.0187 |
| Phenylalanine | Amino Acid | 1.3499 | 0.43286 | 0.0010143 | 2.9938 |
| Aconitate | TCA Cycle | 1.4528 | 0.53887 | 0.0011904 | 2.9243 |
| IMP | Nucleotide/Purine metabolism | 1.4803 | 0.56588 | 0.0015179 | 2.8187 |
| Pantothenic acid | Vitamin/CoA | 1.7378 | 0.79725 | 0.0015518 | 2.8092 |
| NADP | Nicotinate and nicotinamide metabolism | 2.0291 | 1.0209 | 0.002006 | 2.6977 |
| Trigonelline | Nicotinate and nicotinamide metabolism | 0.72696 | -0.46005 | 0.0024475 | 2.6113 |
| Oxidized glutathione | Amino acid metabolism | 1.6918 | 0.75859 | 0.0026826 | 2.5714 |
| CDP | Nucleotide/Pyrimidine metabolism | 1.8668 | 0.90055 | 0.0029955 | 2.5235 |
| N1-Methylnicotinamide | NAD Metabolism | 1.6302 | 0.70508 | 0.0030881 | 2.5103 |
| NADPH | Nicotinate and nicotinamide metabolism | 1.8407 | 0.88023 | 0.0040566 | 2.3918 |
| Pyroglutamic acid | Amino Acid | 1.6497 | 0.7222 | 0.0060116 | 2.221 |
| Tryptophan | Amino Acid | 1.5738 | 0.65422 | 0.0064967 | 2.1873 |
| GDP | Nucleotide/Purine metabolism | 1.8341 | 0.87507 | 0.0078247 | 2.1065 |
| G3P | Glycolysis | 1.322 | 0.40268 | 0.0078706 | 2.104 |
| Cysteine | Amino Acid | 0.60178 | -0.73268 | 0.008608 | 2.0651 |
| UDP | Nucleotide/Pyrimidine metabolism | 1.4385 | 0.52452 | 0.0088668 | 2.0522 |
| Myristoylcarnitine | Lipid metabolism/Fatty Acid metabolism | 2.0372 | 1.0266 | 0.010333 | 1.9858 |
| Xanthine | Nucleotide | 1.5758 | 0.65605 | 0.010804 | 1.9664 |
| Tyrosine | Amino Acid | 0.64328 | -0.63648 | 0.01205 | 1.919 |
| Nicotinamide Riboside | Nicotinate and nicotinamide metabolism | 1.5466 | 0.62907 | 0.016849 | 1.7734 |
| D-2-hydroxyglutarate | TCA cycle | 1.3012 | 0.37988 | 0.018187 | 1.7402 |
| UTP | Nucleotide/pyrimidine | 1.7443 | 0.80261 | 0.018249 | 1.7388 |
| ADP | Nucleotide/Purine metabolism | 1.4149 | 0.50071 | 0.019531 | 1.7093 |
| Methionine | Amino Acid | 0.69156 | -0.53208 | 0.019937 | 1.7003 |
| FAD | Nucleotide | 1.541 | 0.62388 | 0.020585 | 1.6864 |
| Palmitoyl-CoA | Lipid metabolism/Fatty Acid metabolism | 1.9143 | 0.93683 | 0.021527 | 1.667 |
| Lysine | Amino Acid | 0.76526 | -0.38599 | 0.030136 | 1.5209 |
| Palmitate | Lipid metabolism/Fatty Acids | 0.53632 | -0.89884 | 0.036594 | 1.4366 |
| Stearic acid | Lipid metabolism/Fatty Acid | 0.6938 | -0.5274 | 0.038629 | 1.4131 |
| Kynurenine | Amino Acid metabolism/Trp | 1.6981 | 0.76389 | 0.039618 | 1.4021 |
| Guanine | Nucleotide/Purine metabolism | 1.6336 | 0.70809 | 0.049879 | 1.3021 |

**Supplemental Table S4**. Changed retinal metabolites in different metabolic pathways in fed vs. fasted female mice by Volcano plots.

| **Metabolite** | **Pathway** | **FC** | **log2(FC)** | **P Value** | **-log10(p)** |
| --- | --- | --- | --- | --- | --- |
| 3-Hydroxybutyric acid | Lipid/Ketone bodies | 6.1512 | 2.6209 | 6.38E-09 | 8.1955 |
| Trigonelline | Nicotinate and nicotinamide metabolism | 0.43467 | -1.202 | 1.80E-05 | 4.7458 |
| Ascorbic acid | Ascorbate and aldarate metabolism | 1.3844 | 0.46922 | 0.00054799 | 3.2612 |
| Phosphocreatine | Amino Acid/Pro and Arg | 0.76063 | -0.39473 | 0.00071183 | 3.1476 |
| Hexanoylcarnitine | Acylcarnitines/Fatty Acid metabolism | 2.2721 | 1.184 | 0.0017945 | 2.7461 |
| Palmitoylcarnitine | Acylcarnitines/Fatty Acid metabolism | 2.8586 | 1.5153 | 0.0018785 | 2.7262 |
| L-Arginine | Amino Acid/Urea cycle | 0.5904 | -0.76024 | 0.0024084 | 2.6183 |
| Stearoylcarnitine | Acylcarnitines/Fatty Acid metabolism | 2.4699 | 1.3044 | 0.0028844 | 2.5399 |
| Myristoylcarnitine | Lipid metabolism/Fatty Acid metabolism | 2.5431 | 1.3466 | 0.0033302 | 2.4775 |
| Arginosuccinate | Urea Cycle | 0.62751 | -0.67229 | 0.0042092 | 2.3758 |
| Aspartic acid | Amino Acid | 0.70485 | -0.50461 | 0.0048872 | 2.3109 |
| Serine | Amino Acid | 0.53553 | -0.90095 | 0.0066322 | 2.1783 |
| Alanine | Amino Acid | 0.75038 | -0.41431 | 0.0090249 | 2.0446 |
| Pantothenic acid | Vitamin/CoA | 1.3511 | 0.43416 | 0.01189 | 1.9248 |
| Propionylcarnitine | Acylcarnitine/Fatty Acid metabolism | 1.9806 | 0.98592 | 0.012951 | 1.8877 |
| Carnitine | Amino Acid metabolism/Lys | 0.74072 | -0.43301 | 0.015299 | 1.8153 |
| Aminoadipic acid | Amino Acid metabolism/Ly | 0.70052 | -0.51351 | 0.020473 | 1.6888 |
| Threonine | Amino Acid | 0.69972 | -0.51514 | 0.024436 | 1.612 |
| Methionine | Amino Acid | 0.55488 | -0.84975 | 0.028497 | 1.5452 |
| L-Homoserine | Amino Acid/Thr, Met, Asp | 0.71292 | -0.48818 | 0.029511 | 1.53 |
| Kynurenine | Amino Acid metabolism/Trp | 1.457 | 0.54301 | 0.041153 | 1.3856 |

**Supplemental Table S5**. Changed RPE metabolites in different metabolic pathways in the fed state by Volcano plots.

| **Metabolite** | **Pathway** | **FC** | **log2(FC)** | **P Value** | **-log10(p)** |
| --- | --- | --- | --- | --- | --- |
| Succinate | TCA Cycle | 0.28256 | -1.8233 | 5.76E-06 | 5.2395 |
| Beta-Alanine | Amino Acid | 0.50734 | -0.97897 | 0.0001627 | 3.7887 |
| Taurine | Amino Acid | 0.61686 | -0.69698 | 0.0026604 | 2.5751 |
| 4-Hydroxyproline | Amino Acid metabolism/Pro, Arg | 0.61081 | -0.71121 | 0.0028086 | 2.5515 |
| Acetyl-CoA | TCA Cycle/Fatty acid metabolism | 0.58175 | -0.78152 | 0.0046309 | 2.3343 |
| AcetylCholine | Lipid/phospholipid, | 0.30612 | -1.7078 | 0.0075427 | 2.1225 |
| Hypotaurine | Taurine metabolism | 0.51827 | -0.94823 | 0.0078318 | 2.1061 |
| Hypoxanthine | Nucleotide/Purine metabolism | 0.71865 | -0.47664 | 0.011852 | 1.9262 |
| ATP | Nucleotide | 0.50062 | -0.9982 | 0.012307 | 1.9098 |
| NAD | Nicotinate and nicotinamide metabolism | 0.69381 | -0.52739 | 0.015164 | 1.8192 |
| Citrate | TCA Cycle | 0.70483 | -0.50466 | 0.018697 | 1.7282 |
| Creatinine | Amino Acids/Arg, gly | 0.56817 | -0.8156 | 0.022438 | 1.649 |
| Pantothenic acid | Vitamin/CoA | 1.4528 | 0.53888 | 0.026994 | 1.5687 |
| Histamine | Amino acid metabolism | 0.66588 | -0.58666 | 0.028587 | 1.5438 |
| Guanine | Nucleotide/Purine metabolism | 0.66912 | -0.57967 | 0.031756 | 1.4982 |
| Palmitoylcarnitine | Acylcarnitines/Fatty Acid metabolism | 0.72661 | -0.46075 | 0.032625 | 1.4865 |
| Propionylcarnitine | Acylcarnitine/Fatty Acid metabolism | 0.72521 | -0.46353 | 0.034629 | 1.4606 |
| D-Ribulose 5-phosphate | Glycolysis/PPP | 0.67922 | -0.55806 | 0.040824 | 1.3891 |
| Carnosine | Amino Acids/Histidine | 0.57018 | -0.81052 | 0.048639 | 1.313 |

**Supplemental Table S6**. Changed RPE metabolites in different metabolic pathways in fed vs. fasted male mice by Volcano plots.

| **Metabolite** | **Pathway** | **FC** | **log2(FC)** | **P Value** | **-log10(p)** |
| --- | --- | --- | --- | --- | --- |
| 3-Hydroxybutyric acid | Lipid/Ketone bodies | 7.6816 | 2.9414 | 6.65E-05 | 4.1774 |
| Trigonelline | Nicotinate and nicotinamide metabolism | 0.44784 | -1.1589 | 0.0002962 | 3.5284 |
| Palmitoylcarnitine | Acylcarnitines/Fatty Acid metabolism | 2.333 | 1.2222 | 0.0004746 | 3.3237 |
| AcetylCarnitine | Lipid/phospholipid | 1.7073 | 0.7717 | 0.0005647 | 3.2482 |
| Isoleucine | Amino Acid | 1.5756 | 0.65586 | 0.000902 | 3.0448 |
| Hexanoylcarnitine | Acylcarnitines/Fatty Acid metabolism | 2.3603 | 1.239 | 0.0013745 | 2.8618 |
| Serine | Amino Acid | 0.74735 | -0.42013 | 0.0015128 | 2.8202 |
| Oxidized glutathione | Amino acid metabolism | 0.70137 | -0.51175 | 0.002598 | 2.5854 |
| Myristoylcarnitine | Lipid metabolism/Fatty Acid metabolism | 2.608 | 1.383 | 0.0027566 | 2.5596 |
| 2-Methylbutyroylcarnitine | AcylCarnitine/Amino acid metabolism | 0.56719 | -0.8181 | 0.005489 | 2.2605 |
| Hypoxanthine | Nucleotide/Purine metabolism | 0.74717 | -0.42049 | 0.0066116 | 2.1797 |
| Hypotaurine | Taurine metabolism | 0.72661 | -0.46076 | 0.0077346 | 2.1116 |
| Aspartic acid | Amino Acid | 0.675 | -0.56704 | 0.0081778 | 2.0874 |
| Tyrosine | Amino Acid | 0.66191 | -0.5953 | 0.0082668 | 2.0827 |
| Malate | TCA Cycle | 1.348 | 0.43078 | 0.0098496 | 2.0066 |
| Urea | Urea Cycle | 1.4937 | 0.57887 | 0.013164 | 1.8806 |
| Aminoadipic acid | Amino Acid metabolism/Lysine | 1.9418 | 0.95738 | 0.014115 | 1.8503 |
| Propionylcarnitine | Acylcarnitine/Fatty Acid metabolism | 0.69463 | -0.52568 | 0.020135 | 1.696 |
| Cytidine | Nucleotide | 1.4469 | 0.53297 | 0.020744 | 1.6831 |
| Stearoylcarnitine | Acylcarnitines/Fatty Acid metabolism | 1.6847 | 0.75249 | 0.023764 | 1.6241 |
| Pyruvate | Glycolysis | 1.3479 | 0.43076 | 0.027049 | 1.5678 |
| UDP | Nucleotide/Pyrimidine metabolism | 0.75831 | -0.39913 | 0.027087 | 1.5672 |

**Supplemental Table S7**. Changed RPE metabolites in different metabolic pathways in fed vs. fasted female mice by Volcano plots.

| **Metabolite** | **Pathway** | **FC** | **log2(FC)** | **P Value** | **-log10(p)** |
| --- | --- | --- | --- | --- | --- |
| Trigonelline | Nicotinate and nicotinamide metabolism | 0.37264 | -1.4241 | 3.46E-05 | 4.4613 |
| Hexanoylcarnitine | Acylcarnitines/Fatty Acid metabolism | 2.023 | 1.0165 | 0.0002045 | 3.6892 |
| 3-Hydroxybutyric acid | Lipid/Ketone bodies | 7.6035 | 2.9267 | 0.0003618 | 3.4415 |
| Stearoylcarnitine | Acylcarnitines/Fatty Acid metabolism | 2.2963 | 1.1993 | 0.0005578 | 3.2535 |
| Myristoylcarnitine | Lipid metabolism/Fatty Acid metabolism | 3.2121 | 1.6835 | 0.0009731 | 3.0119 |
| AcetylCarnitine | Lipid metabolism | 1.6574 | 0.72896 | 0.0012461 | 2.9044 |
| Isoleucine | Amino Acid | 1.554 | 0.63598 | 0.0017928 | 2.7465 |
| Palmitoylcarnitine | Acylcarnitines/Fatty Acid metabolism | 2.8697 | 1.5209 | 0.002428 | 2.6147 |
| Aspartic acid | Amino Acid | 0.61598 | -0.69905 | 0.0064289 | 2.1919 |
| Acetyl-CoA | TCA Cycle/Fatty acid metabolism | 2.1671 | 1.1158 | 0.006723 | 2.1724 |
| Serine | Amino Acid | 0.62834 | -0.67038 | 0.0072295 | 2.1409 |
| 2-Methylbutyroylcarnitine | AcylCarnitine/Amino acid metabolism | 0.49054 | -1.0276 | 0.011665 | 1.9331 |
| Isocitrate | TCA Cycle | 1.5283 | 0.6119 | 0.013093 | 1.883 |
| Citrate | TCA Cycle | 1.632 | 0.70663 | 0.016346 | 1.7866 |
| Malate | TCA Cycle | 1.3829 | 0.46766 | 0.02038 | 1.6908 |
| Creatinine | Amino Acids/Arg, gly | 1.3956 | 0.4809 | 0.024285 | 1.6147 |
| Glutamic acid | Amino Acid | 0.73231 | -0.44947 | 0.027103 | 1.567 |
| L-Arginine | Amino Acid/Urea cycle | 0.65482 | -0.61083 | 0.03111 | 1.5071 |
| Phosphocreatine | Amino Acid/pro and arginine | 1.3621 | 0.44585 | 0.031335 | 1.504 |
| Carnosine | Amino Acids/Histidine | 1.4727 | 0.55848 | 0.045295 | 1.3439 |
| Pantothenic acid | Vitamin/CoA | 1.3527 | 0.43579 | 0.047715 | 1.3213 |
| Cytidine | Nucleotide | 1.3723 | 0.45659 | 0.048568 | 1.3136 |

**Supplemental Table S8**. Changed lens metabolites in different metabolic pathways in the fed state by Volcano plots.

| **Metabolite** | **Pathway** | **FC** | **log2(FC)** | **P Value** | **-log10(p)** |
| --- | --- | --- | --- | --- | --- |
| 4-Hydroxyproline | Amino Acid metabolism/Pro, Arg | 0.42469 | -1.2355 | 1.69E-07 | 6.7712 |
| Hypotaurine | Taurine metabolism | 0.5417 | -0.88443 | 1.61E-05 | 4.7944 |
| Ascorbic acid | Ascorbate and aldarate metabolism | 0.12633 | -2.9847 | 3.80E-05 | 4.4204 |
| Glutathione | Amino acid metabolism | 0.25927 | -1.9475 | 8.57E-05 | 4.0668 |
| UDP-Glucosamine | Nucleotide sugar | 0.4611 | -1.1169 | 0.000487 | 3.3125 |
| Urate | Nucleotide/Purine metabolism | 0.55914 | -0.83872 | 0.0006381 | 3.1951 |
| IMP | Nucleotide/Purine metabolism | 0.68324 | -0.54953 | 0.0007074 | 3.1503 |
| Propionylcarnitine | Acylcarnitine/Fatty Acid metabolism | 0.5577 | -0.84245 | 0.000992 | 3.0035 |
| NADH | Nicotinate and nicotinamide metabolism | 0.4921 | -1.023 | 0.001009 | 2.9961 |
| AMP | Nucleotide | 0.67795 | -0.56074 | 0.0011322 | 2.9461 |
| UDP | Nucleotide/ Pyrimidine metabolism | 0.33224 | -1.5897 | 0.0017076 | 2.7676 |
| CDP | Nucleotide/ Pyrimidine metabolism | 0.3354 | -1.576 | 0.001726 | 2.7629 |
| Kynurenine | Amino Acid metabolism/Trp | 1.5 | 0.58496 | 0.0021433 | 2.6689 |
| D-Ribulose 5-phosphate | Glycolysis/PPP | 0.697 | -0.52077 | 0.0023475 | 2.6294 |
| ADP | Nucleotide/Purine metabolism | 0.41638 | -1.264 | 0.0025508 | 2.5933 |
| GDP | Nucleotide/Purine metabolism | 0.37867 | -1.401 | 0.0041162 | 2.3855 |
| IPP | mevalonate pathway | 0.67784 | -0.56097 | 0.0046131 | 2.336 |
| Butyrylcarnitine | Lipid metabolism/Fatty Acid metabolism | 0.68725 | -0.54109 | 0.0051714 | 2.2864 |
| N-Acetyl-L-aspartic acid | Amino Acid metabolism/Ala, Glu, Asp | 0.65407 | -0.61249 | 0.0053861 | 2.2687 |
| Isobutyrylcarnitine | Lipid metabolism/Fatty Acid metabolism | 0.70134 | -0.51182 | 0.0063538 | 2.197 |
| Cystine | Amino Acid | 1.6243 | 0.69978 | 0.0066549 | 2.1769 |
| Hexanoylcarnitine | Acylcarnitines/Fatty Acid metabolism | 0.6721 | -0.57325 | 0.0069427 | 2.1585 |
| Palmitoylcarnitine | Acylcarnitines/Fatty Acid metabolism | 0.73883 | -0.43669 | 0.0073081 | 2.1362 |
| Erythritol | Sugar | 0.76338 | -0.38952 | 0.0092814 | 2.0324 |
| 2-Methylbutyroylcarnitine | AcylCarnitine/Amino acid metabolism | 0.72039 | -0.47316 | 0.0099399 | 2.0026 |
| Phosphocreatine | Amino Acid/pro and arginine | 0.49136 | -1.0251 | 0.010897 | 1.9627 |
| Myristoylcarnitine | Lipid metabolism/Fatty Acid metabolism | 0.61277 | -0.70658 | 0.012593 | 1.8999 |
| Glucose | Glycolysis/sugar | 1.4773 | 0.56298 | 0.012673 | 1.8971 |
| Acetylglutamic acid | Amino Acid metabolism/Proline/Arginine | 0.76847 | -0.37993 | 0.012866 | 1.8906 |
| Nicotinamide Riboside | Nicotinate and nicotinamide metabolism | 0.66388 | -0.59102 | 0.013104 | 1.8826 |
| GMP | Nucleotide/Purine metabolism | 0.629 | -0.66887 | 0.013846 | 1.8587 |
| Glucose 6-phosphate | Glycolysis/PPP | 0.71809 | -0.47776 | 0.020177 | 1.6951 |
| AcetylCholine | Lipid/phospholipid, ligang | 0.61312 | -0.70577 | 0.024482 | 1.6111 |
| Adenine | Nucleotide/Purine metabolism | 2.2215 | 1.1515 | 0.025506 | 1.5934 |
| Fumarate | TCA Cycle | 0.62599 | -0.67578 | 0.025734 | 1.5895 |
| Arginosuccinate | Urea Cycle | 0.67647 | -0.5639 | 0.028288 | 1.5484 |
| UTP | Nucleotide/pyrimidine | 0.48947 | -1.0307 | 0.03114 | 1.5067 |
| Aconitate | TCA Cycle | 0.61004 | -0.71301 | 0.032692 | 1.4856 |
| Uracil | Nucleotide/ Pyrimidine metabolism | 0.73613 | -0.44196 | 0.033105 | 1.4801 |
| Tryptophan | Amino Acid | 1.3055 | 0.38462 | 0.036312 | 1.44 |
| Isocitrate | TCA Cycle | 0.56684 | -0.81899 | 0.037813 | 1.4224 |
| Malate | TCA Cycle | 0.67148 | -0.57457 | 0.039126 | 1.4075 |
| Citrate | TCA Cycle | 0.6029 | -0.73002 | 0.041385 | 1.3832 |
| a-Ketoglutarate | TCA Cycle | 0.60481 | -0.72544 | 0.042366 | 1.373 |
| 3-Phosphoglyceric acid | Glycolysis | 0.55339 | -0.85364 | 0.048102 | 1.3178 |

**Supplemental Table S9**. Changed lens metabolites in different metabolic pathways in fed vs. fasted male mice by Volcano plots.

| **Metabolite** | **Pathway** | **FC** | **log2(FC)** | **P Value** | **-log10(p)** |
| --- | --- | --- | --- | --- | --- |
| 4-Hydroxyproline | Amino Acid metabolism/Proline/Arginine | 0.45368 | -1.1403 | 1.06E-07 | 6.9736 |
| NADH | Nicotinate and nicotinamide metabolism | 0.52409 | -0.9321 | 5.55E-07 | 6.2555 |
| ADP | Nucleotide/Purine metabolism | 0.37578 | -1.412 | 6.89E-07 | 6.1619 |
| UDP | Nucleotide/Pyrimidine metabolism | 0.29544 | -1.7591 | 3.26E-06 | 5.4863 |
| CDP | Nucleotide/Pyrimidine metabolism | 0.26815 | -1.8989 | 3.86E-06 | 5.4129 |
| IMP | Nucleotide/Purine metabolism | 0.70389 | -0.50658 | 6.97E-06 | 5.1568 |
| AMP | Nucleotide | 0.68836 | -0.53877 | 8.22E-06 | 5.0853 |
| Glutathione | Amino acid metabolism | 0.28772 | -1.7973 | 1.61E-05 | 4.7922 |
| GDP | Nucleotide/Purine metabolism | 0.3004 | -1.735 | 2.40E-05 | 4.6207 |
| Ascorbic acid | Ascorbate and aldarate metabolism | 0.13111 | -2.9312 | 3.13E-05 | 4.5041 |
| Hypotaurine | Taurine metabolism | 0.6706 | -0.57648 | 6.82E-05 | 4.1663 |
| GMP | Nucleotide/Purine metabolism | 0.64486 | -0.63293 | 9.42E-05 | 4.026 |
| ATP | Nucleotide | 0.40133 | -1.3171 | 0.00024 | 3.6194 |
| Adenine | Nucleotide/Purine metabolism | 2.2561 | 1.1739 | 0.00043 | 3.3668 |
| IPP | mevalonate pathway | 0.74224 | -0.43004 | 0.000651 | 3.1861 |
| Aminoadipic acid | Amino Acid metabolism/Lysine | 0.63879 | -0.64658 | 0.000893 | 3.0492 |
| XMP | Nucleotide/Purine metabolism | 0.6833 | -0.54941 | 0.001574 | 2.803 |
| Glucose | Glycolysis/sugar | 2.0243 | 1.0174 | 0.00194 | 2.7122 |
| Nicotinamide riboside | Nicotinate and nicotinamide metabolism | 0.71199 | -0.49007 | 0.002375 | 2.6244 |
| Pantothenic acid | Vitamin/CoA | 1.5388 | 0.62182 | 0.003703 | 2.4314 |
| Thiamine | Vitamins | 0.44959 | -1.1533 | 0.003934 | 2.4052 |
| Cystine | Amino Acid | 2.3494 | 1.2323 | 0.004484 | 2.3484 |
| 3-Phosphoglyceric acid | Glycolysis | 0.61357 | -0.7047 | 0.009814 | 2.0082 |
| Glycerate | Glycolysis | 2.0418 | 1.0298 | 0.015175 | 1.8189 |
| UTP | Nucleotide/pyrimidine | 0.5478 | -0.86828 | 0.015874 | 1.7993 |
| UDP-Glucosamine | Nucleotide sugar | 0.72676 | -0.46045 | 0.016041 | 1.7948 |
| Uracil | Nucleotide/Pyrimidine metabolism | 1.9515 | 0.96461 | 0.016134 | 1.7923 |
| NAD | Nicotinate and nicotinamide metabolism | 0.6555 | -0.60933 | 0.028197 | 1.5498 |

**Supplemental Table S10**. Changed lens metabolites in different metabolic pathways in fed vs. fasted female mice by Volcano plots.

| **Metabolite** | **Pathway** | **FC** | **log2(FC)** | **P Value** | **-log10(p)** |
| --- | --- | --- | --- | --- | --- |
| Methionine | Amino Acid | 0.59031 | -0.76045 | 1.41E-10 | 9.8516 |
| Tyrosine | Amino Acid | 0.61037 | -0.71224 | 1.08E-08 | 7.9659 |
| Glucose | Glycolysis/sugar | 0.3747 | -1.4162 | 1.55E-08 | 7.81 |
| Alanine | Amino Acid | 0.67223 | -0.57297 | 5.37E-08 | 7.2701 |
| Proline | Amino Acid | 0.74338 | -0.42782 | 6.30E-07 | 6.201 |
| Kynurenine | Amino Acid metabolism/Trp | 2.04 | 1.0286 | 7.50E-07 | 6.1249 |
| Serine | Amino Acid | 0.73233 | -0.44943 | 1.05E-06 | 5.9786 |
| Aminoadipic acid | Amino Acid metabolism/Lysine | 2.2399 | 1.1634 | 2.67E-06 | 5.5729 |
| 3-Hydroxybutyric acid | Lipid/Ketone bodies | 11.264 | 3.4937 | 3.70E-06 | 5.4319 |
| L-Arginine | Amino Acid/Urea cycle | 0.65624 | -0.60771 | 6.15E-05 | 4.211 |
| Trigonelline | Nicotinate and nicotinamide metabolism | 0.68732 | -0.54095 | 0.0001234 | 3.9088 |
| Glycerate | Glycolysis | 0.62468 | -0.6788 | 0.0003331 | 3.4774 |
| Propionylcarnitine | Acylcarnitine/Fatty Acid metabolism | 1.4952 | 0.58031 | 0.0003437 | 3.4638 |
| XMP | Nucleotide/Purine metabolism | 1.3544 | 0.43763 | 0.001212 | 2.9165 |
| Arginosuccinate | Urea Cycle | 0.67479 | -0.56749 | 0.0023894 | 2.6217 |
| Tryptophan | Amino Acid | 1.4758 | 0.56152 | 0.0033357 | 2.4768 |
| GMP | Nucleotide/Purine metabolism | 1.3321 | 0.41369 | 0.0054533 | 2.2633 |
| ATP | Nucleotide | 1.5326 | 0.61596 | 0.015059 | 1.8222 |
| Citrulline | Urea Cycle | 0.76887 | -0.37919 | 0.016817 | 1.7743 |
| Glycine | Amino Acid | 1.6799 | 0.7484 | 0.030133 | 1.521 |

**Supplemental Table S11**. Changed brain metabolites in different metabolic pathways in the fed state by Volcano plots.

| **Metabolite** | **Pathway** | **FC** | **log2(FC)** | **P Value** | **-log10(p)** |
| --- | --- | --- | --- | --- | --- |
| 3-Phosphoglyceric acid | Glycolysis | 1.6517 | 0.72392 | 0.017088 | 1.7673 |
| cAMP | Nucleotide/Purine metabolism | 1.9955 | 0.99672 | 0.031295 | 1.5045 |
| Glucose | Glycolysis/sugar | 2.1654 | 1.1147 | 0.028782 | 1.5409 |
| Glucose 1-phosphate | Glycolysis | 1.3453 | 0.42793 | 0.01782 | 1.7491 |
| Glucose 6-phosphate | Glycolysis/PPP | 1.4109 | 0.49657 | 0.024843 | 1.6048 |
| Glycerate | Glycolysis | 2.4906 | 1.3165 | 0.021429 | 1.669 |
| Hypotaurine | Taurine metabolism | 0.43133 | -1.2131 | 0.010304 | 1.987 |
| Hypoxanthine | Nucleotide/Purine metabolism | 0.68879 | -0.53787 | 0.001358 | 2.867 |
| Kynurenine | Amino Acid metabolism/Trp | 1.3636 | 0.44747 | 0.047157 | 1.3265 |
| NAD | Nicotinate and nicotinamide metabolism | 0.67971 | -0.55701 | 0.024357 | 1.6134 |
| Pantothenic acid | Vitamin/CoA | 1.3712 | 0.45546 | 0.000156 | 3.8081 |
| PEP | Glycolysis | 1.4706 | 0.55639 | 0.018721 | 1.7277 |
| Thiamine | Vitamins | 0.59737 | -0.74329 | 0.025238 | 1.5979 |
| Uracil | Nucleotide/Pyrimidine metabolism | 0.74944 | -0.41611 | 0.011571 | 1.9366 |

**Supplemental Table S12**. Changed brain metabolites in different metabolic pathways in the fasted state by Volcano plots.

| **Metabolite** | **Pathway** | **FC** | **log2(FC)** | **P Value** | **-log10(p)** |
| --- | --- | --- | --- | --- | --- |
| Aconitate | TCA Cycle | 1.8474 | 0.88546 | 0.023852 | 1.6225 |
| ADP | Nucleotide/Purine metabolism | 0.7174 | -0.47915 | 0.049835 | 1.3025 |
| ATP | Nucleotide | 0.27504 | -1.8623 | 0.013567 | 1.8675 |
| cGMP | Nucleotide/Purine metabolism | 2.0395 | 1.0282 | 0.023383 | 1.6311 |
| CoA | TCA Cycle/Fatty acid metabolism | 0.4994 | -1.0017 | 0.040616 | 1.3913 |
| Cystathionine | Amino Acids/cys | 1.6675 | 0.73766 | 0.006593 | 2.1809 |
| Hypotaurine | Taurine metabolism | 0.45918 | -1.1229 | 0.049854 | 1.3023 |
| Methionine | Amino Acid | 0.5614 | -0.83289 | 0.023608 | 1.6269 |
| NADP | Nicotinate and nicotinamide metabolism | 0.41096 | -1.2829 | 0.028343 | 1.5475 |
| Nicotinamide riboside | Nicotinate and nicotinamide metabolism | 0.68593 | -0.54387 | 0.045193 | 1.3449 |
| Oxidized glutathione | Amino acid metabolism | 0.45622 | -1.1322 | 0.041305 | 1.384 |
| Pantothenic acid | Vitamin/CoA | 1.474 | 0.55969 | 0.000472 | 3.3258 |
| SAM | Amino Acid/pro and arginine | 0.4397 | -1.1854 | 0.007834 | 2.106 |
| Tyrosine | Amino Acid | 0.5899 | -0.76147 | 0.004907 | 2.3092 |
| Xanthine | Nucleotide | 0.66431 | -0.59007 | 0.007764 | 2.1099 |

**Supplemental Table S13**. Changed brain metabolites in different metabolic pathways in the fasted state by Volcano plots.

| **Metabolite** | **Pathway** | **FC** | **log2(FC)** | **P Value** | **-log10(p)** |
| --- | --- | --- | --- | --- | --- |
| Aconitate | TCA Cycle | 1.8474 | 0.88546 | 0.023852 | 1.6225 |
| ADP | Nucleotide/Purine metabolism | 0.7174 | -0.47915 | 0.049835 | 1.3025 |
| ATP | Nucleotide | 0.27504 | -1.8623 | 0.013567 | 1.8675 |
| cGMP | Nucleotide/Purine metabolism | 2.0395 | 1.0282 | 0.023383 | 1.6311 |
| CoA | TCA Cycle/Fatty acid metabolism | 0.4994 | -1.0017 | 0.040616 | 1.3913 |
| Cystathionine | Amino Acids/cys | 1.6675 | 0.73766 | 0.006593 | 2.1809 |
| Hypotaurine | Taurine metabolism | 0.45918 | -1.1229 | 0.049854 | 1.3023 |
| Methionine | Amino Acid | 0.5614 | -0.83289 | 0.023608 | 1.6269 |
| NADP | Nicotinate and nicotinamide metabolism | 0.41096 | -1.2829 | 0.028343 | 1.5475 |
| Nicotinamide riboside | Nicotinate and nicotinamide metabolism | 0.68593 | -0.54387 | 0.045193 | 1.3449 |
| Oxidized glutathione | Amino acid metabolism | 0.45622 | -1.1322 | 0.041305 | 1.384 |
| Pantothenic acid | Vitamin/CoA | 1.474 | 0.55969 | 0.000472 | 3.3258 |
| SAM | Amino Acid/pro and arginine | 0.4397 | -1.1854 | 0.007834 | 2.106 |
| Tyrosine | Amino Acid | 0.5899 | -0.76147 | 0.004907 | 2.3092 |
| Xanthine | Nucleotide | 0.66431 | -0.59007 | 0.007764 | 2.1099 |

**Supplemental Table S14**. Changed brain metabolites in different metabolic pathways in fed vs. fasted male mice by Volcano plots.

| **Metabolite** | **Pathway** | **FC** | **log2(FC)** | **P Value** | **-log10(p)** |
| --- | --- | --- | --- | --- | --- |
| 2-Methylbutyroylcarnitine | AcylCarnitine/Amino acid metabolism | 1.4493 | 0.53532 | 0.0036333 | 2.4397 |
| Aminoadipic acid | Amino Acid metabolism/Lysine | 1.3228 | 0.40356 | 0.0045172 | 2.3451 |
| Urate | Nucleotide/Purine metabolism | 1.3555 | 0.43885 | 0.0057283 | 2.242 |
| Trigonelline | Nicotinate and nicotinamide metabolism | 0.62827 | -0.67054 | 0.0068483 | 2.1644 |
| 3-Hydroxybutyric acid | Lipid/Ketone bodies | 4.239 | 2.0837 | 0.014167 | 1.8487 |
| N1-Methylnicotinamide | NAD Metabolism | 1.3323 | 0.41389 | 0.019394 | 1.7123 |
| G3P | Glycolysis | 0.60539 | -0.72407 | 0.020246 | 1.6937 |
| NADP | Nicotinate and nicotinamide metabolism | 0.60551 | -0.72378 | 0.027281 | 1.5641 |
| UTP | Nucleotide/pyrimidine | 0.46316 | -1.1104 | 0.034957 | 1.4565 |

**Supplemental Table S15**. Changed brain metabolites in different metabolic pathways in fed vs. fasted female mice by Volcano plots.

| **Metabolite** | **Pathway** | **FC** | **log2(FC)** | **P Value** | **-log10(p)** |
| --- | --- | --- | --- | --- | --- |
| 3-Hydroxybutyric acid | Lipid/Ketone bodies | 3.951 | 1.9822 | 1.17E-07 | 6.9321 |
| Isoleucine | Amino Acid | 1.3255 | 0.40648 | 3.78E-05 | 4.4228 |
| Tyrosine | Amino Acid | 0.47872 | -1.0627 | 0.0002374 | 3.6246 |
| Trigonelline | Nicotinate and nicotinamide metabolism | 0.51904 | -0.94609 | 0.0002443 | 3.6121 |
| 2-Methylbutyroylcarnitine | AcylCarnitine/Amino acid metabolism | 1.398 | 0.48333 | 0.0011989 | 2.9212 |
| Lactate | Glycolysis | 0.76208 | -0.39199 | 0.0013095 | 2.8829 |
| Methionine | Amino Acid | 0.44833 | -1.1574 | 0.0016478 | 2.7831 |
| Kynurenine | Amino Acid metabolism/Trp | 1.6713 | 0.74096 | 0.0029291 | 2.5333 |
| Cystathionine | Amino Acids/cys | 1.7148 | 0.77801 | 0.0033708 | 2.4723 |
| DHAP | Glycolysis | 0.33395 | -1.5823 | 0.005669 | 2.2465 |
| Valine | Amino Acid | 1.5442 | 0.62683 | 0.0057904 | 2.2373 |
| cGMP | Nucleotide/Purine metabolism | 2.1808 | 1.1248 | 0.0075014 | 2.1249 |
| NADH | Nicotinate and nicotinamide metabolism | 0.58112 | -0.7831 | 0.0099077 | 2.004 |
| PEP | Glycolysis | 0.56546 | -0.8225 | 0.010287 | 1.9877 |
| Aconitate | TCA Cycle | 1.8577 | 0.89349 | 0.011694 | 1.932 |
| L-Arginine | Amino Acid/Urea cycle | 0.72455 | -0.46484 | 0.012296 | 1.9102 |
| Oxidized glutathione | Amino acid metabolism | 0.42457 | -1.2359 | 0.012386 | 1.9071 |
| Glucose 1-phosphate | Glycolysis | 0.67156 | -0.57441 | 0.015201 | 1.8181 |
| Lysine | Amino Acid | 0.76242 | -0.39133 | 0.015609 | 1.8066 |
| 3-Phosphoglyceric acid | Glycolysis | 0.54293 | -0.88117 | 0.018662 | 1.729 |
| CDP | Nucleotide/Pyrimidine metabolism | 0.40885 | -1.2903 | 0.019274 | 1.715 |
| UTP | Nucleotide/pyrimidine | 0.32129 | -1.6381 | 0.022425 | 1.6493 |
| ATP | Nucleotide | 0.21247 | -2.2347 | 0.023385 | 1.6311 |
| ADP | Nucleotide/Purine metabolism | 0.67141 | -0.57474 | 0.024548 | 1.61 |
| Butyrylcarnitine | Lipid metabolism/Fatty Acid metabolism | 1.4806 | 0.56619 | 0.02763 | 1.5586 |
| Heptadecanoic acid | Lipid metabolism/Fatty Acid metabolism | 1.3813 | 0.46602 | 0.028489 | 1.5453 |
| NADP | Nicotinate and nicotinamide metabolism | 0.36044 | -1.4722 | 0.029498 | 1.5302 |
| Nicotinamide Riboside | Nicotinate and nicotinamide metabolism | 0.68868 | -0.53809 | 0.03521 | 1.4533 |
| Ornithine | Amino acid/Urea cyle | 0.7114 | -0.49127 | 0.035326 | 1.4519 |
| Isobutyrylcarnitine | Lipid metabolism/Fatty Acid metabolism | 1.4548 | 0.54086 | 0.040416 | 1.3934 |
| Glucose | Glycolysis/sugar | 0.52179 | -0.93845 | 0.041703 | 1.3798 |
| Glucose 6-phosphate | Glycolysis/PPP | 0.71928 | -0.47538 | 0.047523 | 1.3231 |

**Supplemental Table S16.** Changed plasma metabolites in different metabolic pathways in fed vs. fasted male mice by Volcano plots.

| **Metabolite** | **Pathway** | **FC** | **log2(FC)** | **P Value** | **-log10(p)** |
| --- | --- | --- | --- | --- | --- |
| Aminoadipic acid | Amino Acid metabolism/Lysine | 1.9943 | 0.99588 | 0.0009097 | 3.0411 |
| Glucose | Glycolysis/sugar | 0.44742 | -1.1603 | 0.0024337 | 2.6137 |
| Hypotaurine | Taurine metabolism | 0.35 | -1.5146 | 0.0058226 | 2.2349 |
| Trigonelline | Nicotinate and nicotinamide metabolism | 0.20947 | -2.2552 | 0.0065327 | 2.1849 |
| Cis-aconitic acid | TCA Cycle | 0.22744 | -2.1364 | 0.0090333 | 2.0442 |
| IPP | mevalonate pathway | 0.52134 | -0.93971 | 0.009865 | 2.0059 |
| CDP | Nucleotide/Pyrimidine metabolism | 0.48945 | -1.0308 | 0.019984 | 1.6993 |
| Alanine | Amino Acid | 0.64641 | -0.62948 | 0.022802 | 1.642 |
| 3-Hydroxybutyric acid | Lipid/Ketone bodies | 7.4789 | 2.9028 | 0.024978 | 1.6024 |
| Lysine | Amino Acid | 0.59674 | -0.74482 | 0.027688 | 1.5577 |
| Cytidine | Nucleotide | 1.3384 | 0.42056 | 0.04725 | 1.3256 |

**Supplemental Table S17.** Changed plasma metabolites in different metabolic pathways in fed vs. fasted male mice by Volcano plots.

| **Metabolite** | **Pathway** | **FC** | **log2(FC)** | **P Value** | **-log10(p)** |
| --- | --- | --- | --- | --- | --- |
| Trigonelline | Nicotinate and nicotinamide metabolism | 0.15111 | -2.7264 | 8.81E-05 | 4.0548 |
| Glucose | Glycolysis/sugar | 0.50091 | -0.99739 | 0.002212 | 2.6552 |
| Succinate | TCA Cycle | 0.76015 | -0.39564 | 0.0048231 | 2.3167 |
| Cis-aconitic acid | TCA Cycle | 0.44545 | -1.1667 | 0.0066273 | 2.1787 |
| Lactate | Glycolysis | 0.7446 | -0.42546 | 0.018613 | 1.7302 |
| Lysine | Amino Acid | 0.60461 | -0.72593 | 0.020769 | 1.6826 |
| Betaine | Amino Acid/gly,ser, thr metabolism | 0.65289 | -0.61509 | 0.024545 | 1.61 |
| Glutamate | Amino Acid | 0.35916 | -1.4773 | 0.02569 | 1.5902 |
| Aspartate | Amino Acid | 0.40158 | -1.3162 | 0.029626 | 1.5283 |
| 3-Phosphoglyceric acid | Glycolysis | 0.36667 | -1.4475 | 0.029718 | 1.527 |
| Butyrylcarnitine | Lipid metabolism/Fatty Acid metabolism | 0.4669 | -1.0988 | 0.03168 | 1.4992 |
| 3-Hydroxybutyric acid | Lipid/Ketone bodies | 4.5737 | 2.1934 | 0.034554 | 1.4615 |
| Cysteine | Amino Acid | 0.3725 | -1.4247 | 0.039399 | 1.4045 |
| Thiamine | Vitamins | 0.66342 | -0.59201 | 0.04269 | 1.3697 |
| Ornithine | Amino acid/Urea cyle | 0.41608 | -1.2651 | 0.043573 | 1.3608 |
| Isobutyrylcarnitine | Lipid metabolism/Fatty Acid metabolism | 0.51287 | -0.96333 | 0.04642 | 1.3333 |
| Propionylcarnitine | Acylcarnitine/Fatty Acid metabolism | 0.43513 | -1.2005 | 0.046526 | 1.3323 |
